# Two co-occurring dynamics explain the organization of a natural movement repertoire

**DOI:** 10.64898/2026.11.17.767029

**Authors:** Yoav Rubinstein, Maayan Moshkovitz, Shai Tishby Tamari, Stas Tiomkin, Lilach Avitan

## Abstract

Natural movements vary across multiple kinematic features, making it difficult to identify the dynamics shared across movements that generate a behavioral repertoire. Here, we provide a low-dimensional, interpretable generative account of the larval zebrafish hunting repertoire that links tail dynamics to behavioral outcomes. Analysis of thousands of movements spanning classically defined bout types revealed that repertoire diversity arises from continuous combinations of two simultaneously expressed dynamics: a slow, non-oscillatory bending component and a fast, oscillatory undulatory component. An analytical model captured these dynamics with six interpretable parameters and accurately reconstructed movements across the full repertoire and at two developmental stages. Different parameters predicted distinct aspects of movement outcome: maximum tail bend predicted change in heading angle, maximum tail curvature and movement duration predicted distance traveled, and oscillatory asymmetry predicted displacement direction. Relationships among the dynamical parameters constrained the movements and outcomes available to the animal. Key parameters could also be recovered directly from empirical tail kinematics, establishing them as experimentally accessible movement variables rather than model-dependent latent dimensions. Together, these results explain both the diversity and limits of a natural movement repertoire through constrained combinations of co-occurring dynamics.

## Introduction

Animals generate rich repertoires of movements that unfold through coordinated changes in posture over time ^1,2,3,4,5^. Describing this diversity is a central challenge in behavioral neuroscience. Recent advances in tracking methods enable movement to be measured at high spatiotemporal resolution, producing detailed measurements across many features and time points ^6,7^. To make these measurements tractable, they are often reduced to lower-dimensional descriptions. Such descriptions can reveal how movements are organized within a repertoire, identify regions of similarity, and support comparisons across animals, contexts, or conditions ^4,5,8,9,10^. They therefore provide a powerful means of capturing behavioral complexity.

However, such compact descriptions are not always directly interpretable in terms of the underlying movements ^11,12,13^. The axes that organize these descriptions often combine many measured features, so a change along an axis does not necessarily correspond to a specific, interpretable change in movement dynamics. Thus, low-dimensional spaces can reveal the structure of a behavioral repertoire, but they do not necessarily explain which changes in the underlying kinematics account for this structure. Understanding a behavioral repertoire, therefore, requires linking its structure to specific, interpretable dimensions that are grounded in movement dynamics. These dimensions should also account for how variation in movement dynamics relates to changes in the animal’s position and orientation. Such a framework moves beyond describing how the repertoire is organized to explain how movement dynamics generate behavioral diversity.

Larval zebrafish hunting behavior provides a tractable natural behavior for linking the organization of a movement repertoire to interpretable movement dynamics and movement outcome. Here, movement outcome refers to the resulting changes in the animal’s position and orientation. During prey capture, larvae produce a rich repertoire of tail movements that vary across multiple kinematic features ^14,15,16,17,18^. Because the full tail can be tracked at high spatiotemporal resolution, tail dynamics and movement outcome can be measured throughout hunting events ^19,20^. Previous low-dimensional descriptions of zebrafish tail movements have advanced our understanding of behavioral organization by identifying recurrent movement types and by showing that their relative usage changes with behavioral context and development ^5,21,22,23,24,25^. However, the tail-dynamics components underlying the diversity of hunting movements remain unclear. In particular, it is unknown which components vary across the full repertoire, how they combine to shape movement diversity, and how they give rise to different movement outcomes such as distance traveled, displacement direction, and changes in heading angle ^19^.

Here, we show that diverse larval zebrafish hunting movements arise from the simultaneous expression of two coordinated tail dynamics: a slow non-oscillatory bending component and a fast oscillatory undulatory component. We captured these two co-occurring dynamics in a compact analytical model with six interpretable parameters that accurately reconstructed individual movements across the hunting movement repertoire and at two developmental stages. These parameters revealed how specific features of tail dynamics predict movement outcomes: maximum tail bend predicted changes in heading angle, maximum tail curvature and movement duration predicted distance traveled, and oscillatory asymmetry predicted deviations in displacement direction. Importantly, these parameters could also be estimated directly from empirical tail kinematics without fitting the full model to each movement. Using these interpretable parameters, we found that hunting movements vary continuously along these tail-dynamics axes, occupying a constrained region of parameter space. Together, this framework shows how movement dynamics account for diversity across the behavioral repertoire, predict movement outcomes, and provide candidate variables for investigating neural motor control.

## Results

### Tail movements decompose into simultaneous slow bending and fast oscillatory dynamics

We sought to capture tail movements in a low-dimensional manner by treating each movement as a sequence of tail postures evolving over time. We recorded hunting behavior from 30 larval zebrafish (5-7 days post-fertilization, dpf) at 500 Hz (see Methods). Using a custom tracking system, we extracted the fish contour, eye positions, head direction, and tail posture (i.e., the tail midline from the swim bladder to the tip) on a frame-by-frame basis (Figure 1A), generating a time series of tail postures. Hunting events were automatically detected based on eye convergence ^26^ and segmented into individual swim bouts (‘movements’; see Methods, n=7713 movements). For each movement, we quantified the change in fish position and orientation (’movement outcome’ ^19^) (Figure 1B). This dataset allowed us to study tail postural dynamics during hunting and relate them to movement outcomes.

**Figure 1:**
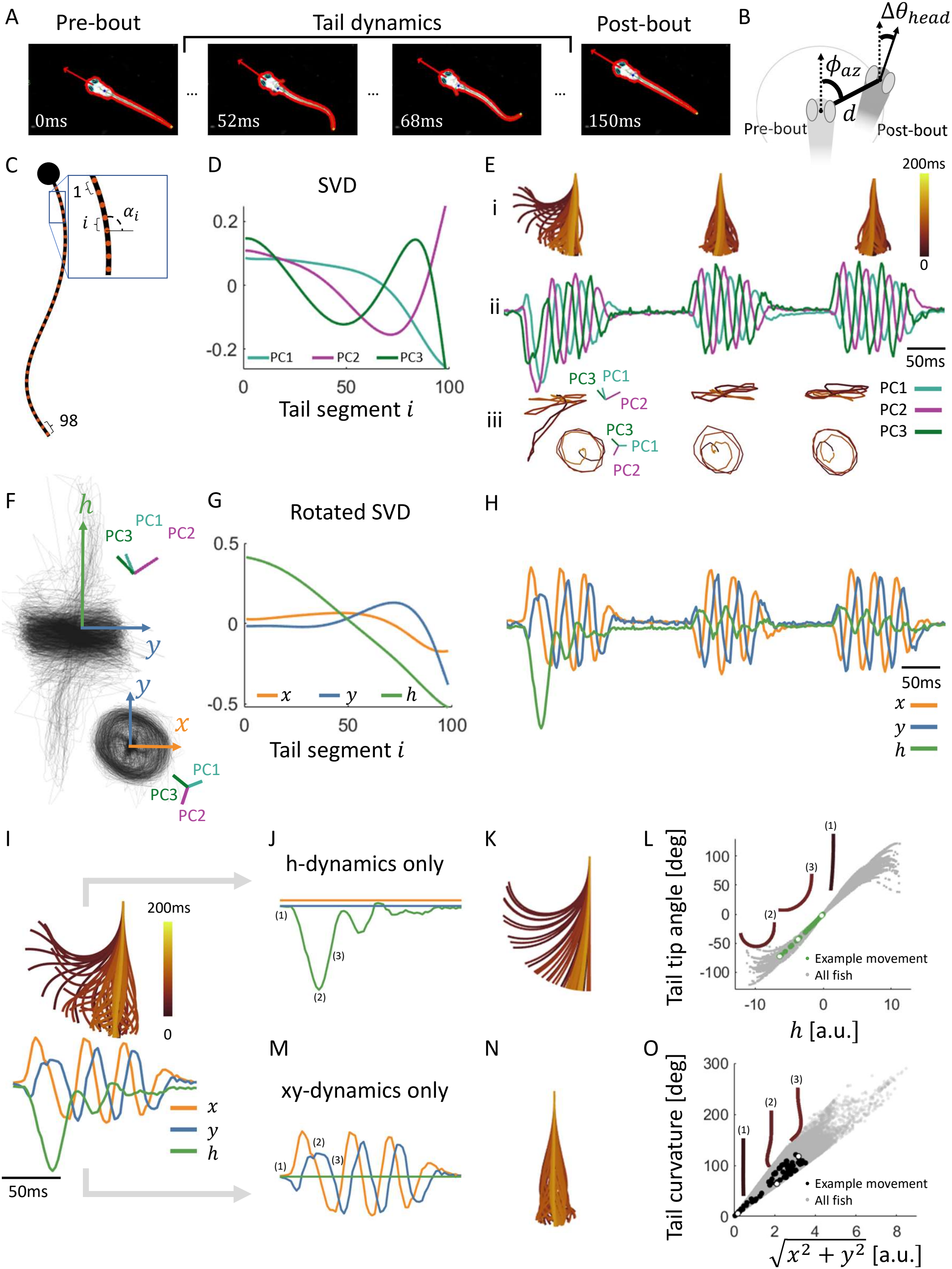
Tail movements decompose into simultaneous slow bending and fast oscillatory dynamics. **A:** Four frames from a representative movement during a hunting event. Fish contour, head direction, fitted eye ellipses, swim bladder, tail midline, and tail tip were tracked and indicated on the frames. **B:** Movement outcome was defined as the change in fish position and orientation resulting from each movement and quantified by three parameters: distance (*d*), change in heading angle (Δ*θ*_*head*_), and azimuthal angle of displacement (*ϕ*_*az*_). Fish position and orientation before and after the movement are indicated by light- and dark-gray fish heads, respectively. Dashed and solid arrows indicate heading direction before and after the movement, respectively. **C:** Tail posture was divided into 98 equally spaced segments and described by a vector of tail segment angles (*α*) relative to the horizontal axis. The black dot indicates the swim bladder. **D:** The first three posture principal components of a representative fish, accounting for 87% of posture variance. **E:** (i) Tail posture dynamics across three consecutive movements over 500 ms of a representative fish. (ii) Time courses of the three principal component coefficients for the same movements. The coefficients varied during each movement and returned toward zero between movements. Across all movements from the representative fish, the coefficients were correlated, with peak cross-correlation across lags of *r*_*PC*1,*PC*2_ = 0.65, *r*_*PC*2,*PC*3_ = 0.74, and *r*_*PC*1,*PC*3_ = 0.63. (iii) The same movements plotted as trajectories in three-dimensional PC space, viewed from above and from the side. The top view reveals rotation at an approximately constant radius, whereas the side view shows the rotational component together with displacement along a third dimension. These trajectories reveal two dominant dynamics: a rotational component present in all three movements, and an additional displacement component, most prominent in the first movement. **F:** Trajectories of all movements from the representative fish in three-dimensional PC space. The movement repertoire occupied a bounded cylinder with a characteristic radius. The principal tail posture component axes (PC1-3) and the transformed cylinder-aligned axes (x, y, and h) are shown. **G:** The posture basis shown in D after rotation into the cylinder-aligned coordinate system (x, y, h). Because the h component varies approximately linearly along the tail, it describes a tail posture that approximates a circular arc. **H:** Coefficients of the three movements shown in E after rotation into the cylinder-aligned coordinates (x, y, h). The x- and y-coefficients showed fast oscillatory dynamics within each movement, whereas h varied more slowly. Across all movements of the representative fish, peak cross-correlations across lags were *r*_*x,y*_ = 0.80, *r*_*x,h*_ = 0.33, *r*_*y,h*_ = 0.31, compare with Eii. **I:** Cylinder-aligned coefficient time series for the first movement shown in E (bottom), together with the corresponding reconstructed tail posture dynamics (top). The time scale bar applies also to the subsequent panels K and N. **J-K:** Setting the xy-dynamics to zero isolates the h-dynamics (J) producing the corresponding unilateral tail bend (K). Three example time points are indicated on the h time series. **L:** The h-dynamics closely tracked tail-tip angle, defined as the angle between the first and the last tail segments. Green dots show this relationship for the example movement in J-K, with white dots indicating the three time points labeled 1-3 in J. Gray dots show data from all postures across all fish (*R*^2^ = 0.99). **M-N:** Setting the h-dynamics to zero isolates the xy-dynamics (M) and produces rhythmic oscillatory undulations (N). The same three time points labeled 1-3 in J are indicated on the xy time series. **O:** The instantaneous radius in the xy-plane 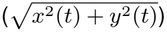 closely tracked tail curvature (see Methods). Black dots show this relationship for the example movement in M-N, with white dots indicating the three time points labeled 1-3 in M. Gray dots show data from all postures across all fish (*R*^2^ = 0.98). All single-fish examples throughout the figure are from the same representative fish.

Zebrafish tail postures are low-dimensional ^22,24^. In our data, the first three principal components of tail posture (’posture components’) explained 83%±4% of the posture variance across fish (mean ± SD), consistent with previous work ^22,24,27^ (Figure 1C-D, see Methods). Their explained variance and shapes were highly consistent across fish (Figure S1A,B). Thus, each instantaneous tail posture during a movement can be characterized by three coefficients, corresponding to the weights of the three principal posture components. A movement, however, is defined by the time course of these coefficients, which oscillate during the movement and return to near zero between movements (Figure 1Ei-ii), highlighting that movement dynamics are higher-dimensional than posture itself.

The time courses of the three posture coefficients showed similar spectral structure and were correlated with one another (Figure S1C,D, Figure 1Eii), suggesting that they were not independent. We therefore asked whether these time courses could be described more compactly. In the three-dimensional PC space, individual movements followed trajectories that rotated at an approximately constant radius within a two-dimensional plane while varying along a third dimension over time (Figure 1Eiii). Comparison with the corresponding tail movements (Movie 1) indicated that the rotational component corresponded to tail undulations, whereas displacement along the third dimension reflected leftward or rightward tail bending. When trajectories from all movements were overlaid, they occupied a bounded cylindrical structure with a characteristic radius (Figure 1F), suggesting that the repertoire is constrained by this geometry and organized by two dominant dynamics: rotation around the cylinder and displacement along its height.

To separate and quantify these two dynamics, we linearly transformed the first three posture components (PC1-PC3) into a cylinder-aligned coordinate system, with one axis along the cylinder height (h) and two spanning its cross-sectional plane (x, y) (Figure 1F,G, Figure S1G-H, see Methods). In this coordinate system, the x- and y-dynamics were tightly coupled to one another but only weakly coupled to h (Figure 1H compared to Figure 1Eii). Consistent with this separation, x and y showed similar coefficient distributions and greater high-frequency spectral power, whereas h showed a different coefficient distribution and was dominated by lower frequencies (Figure S1E-F). The same cylindrical geometry and frequency separation were observed across fish (Figure S1I-L). Although the separation between h-and xy-dynamics was already evident after rotation (Figure 1H; Figure S1K,L), h retained a small residual xy-linked component; removing this component further reduced high-frequency content in h and decreased its coupling to the xy-dynamics (Figure S2; see Methods). Thus, tail movements are compactly described by fast oscillatory (xy) and slow non-oscillatory (h) dynamics.

Given these two dynamics, we next asked how each impacts tail posture. Because tail posture at each time point can be reconstructed from the combined h- and xy-dynamics, we isolated their contributions by reconstructing movements with one dynamic set to zero. Using a representative turn movement as an illustration (Figure 1I), the h-only reconstruction captured a slow unilateral bend and closely tracked the tail-tip angle over time (Figure 1J-L). In contrast, the xy-only reconstruction captured rhythmic tail undulations. The instantaneous radius in the xy plane tracked tail posture curvature, defined as the sum of absolute inter-segment angles^28^ (Figure 1M-O; see Methods). These relationships held robustly across movements and fish (Figure 1L,O). Thus, tail movements decompose into two simultaneously occurring, interpretable dynamics: h-dynamics capture tail bend direction and magnitude, whereas xy-dynamics capture oscillatory alternation between left- and right-curved postures.

### A six-parameter dynamical model reconstructs movements across the hunting movement repertoire

Having identified these two dynamics, we next asked whether their time courses across individual movements could be captured by a compact set of interpretable parameters. In particular, we sought to determine which features of each dynamics must be specified, and how many parameters are sufficient to reproduce the observed diversity of movements. We therefore developed an analytical model that generates the h- and xy-dynamics from these parameters and reconstructs the corresponding tail movements. After removing the xy-linked component, the h-dynamics had a simple time course: it reached an extreme and then returned to baseline, where this extreme corresponded to maximal tail bend (Figure 2A). Accordingly, the h-dynamics were summarized by two interpretable parameters: maximum tail bend (’max tail bend’) and time to maximal bend (’time to bend’). The full fitting model also included two parameters specifying the initial conditions of the movement: its timing relative to the detected movement onset and the small initial offset from straight tail posture at that time (Figure S3A; see Methods). These parameters specify the state of the tail at movement onset rather than the movement dynamics themselves. Thus, the h-dynamics were specified by maximum tail bend and time to bend.

**Figure 2:**
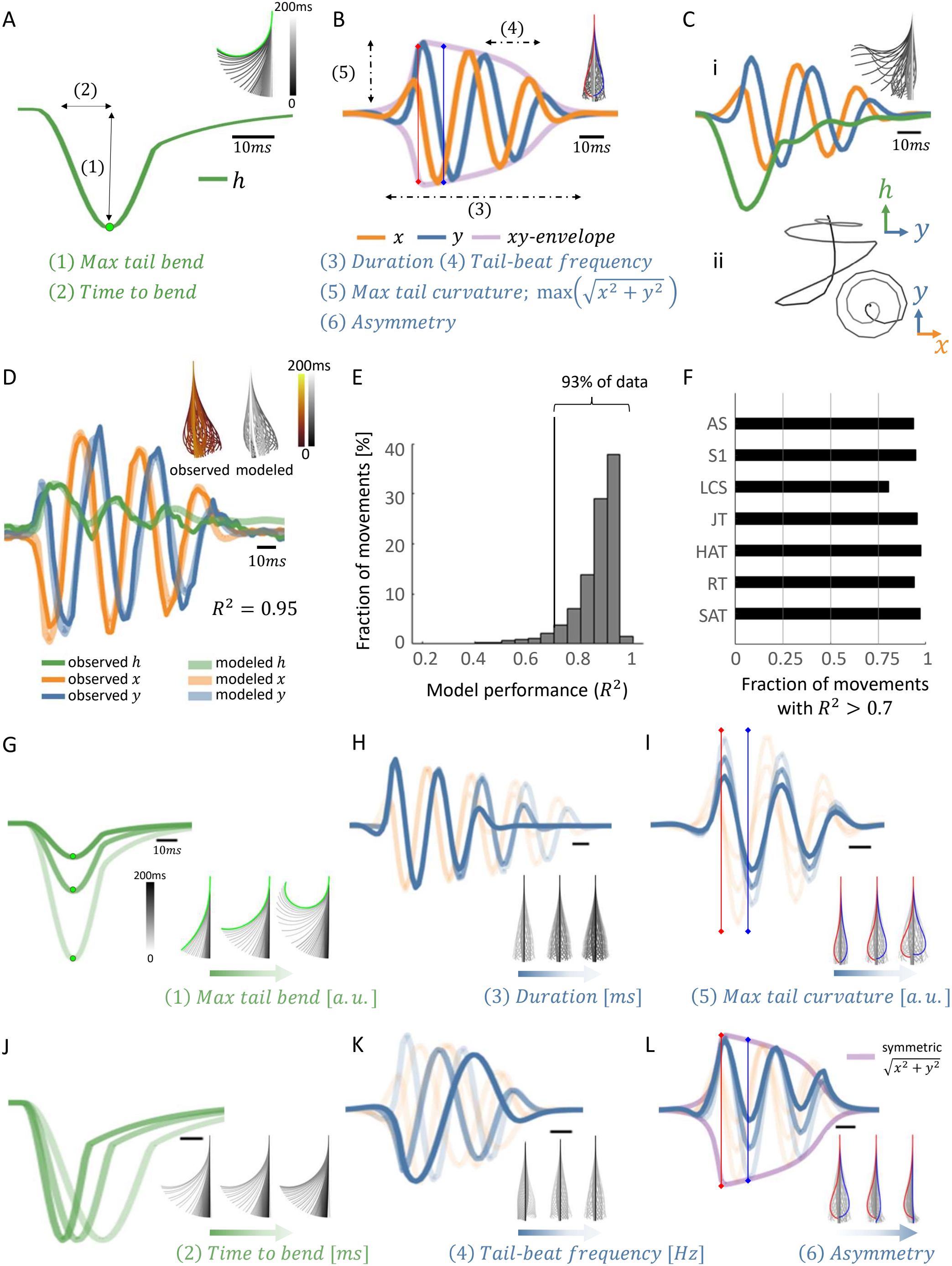
A six-parameter dynamical model reconstructs the hunting movement repertoire. **A:** Left: example h-dynamics, with the green dot marking the extreme value of the dynamics, corresponding to maximal tail bend occurring at the indicated time to bend. Right: tail postures reconstructed from these dynamics, with the green posture indicating the same time point. **B:** Left: the xy-dynamics were captured by an Izhikevich-inspired transient oscillator governed by four interpretable parameters: duration, tail-beat frequency, maximum tail curvature, and asymmetry. The purple curve shows the oscillation envelope, whose peak defines the maximum tail curvature. Red and blue vertical lines mark opposite phases of the oscillation, corresponding to the red and blue maximal leftward and rightward-curved reconstructed tail postures shown on the right. In symmetric movements, oscillations reached similar positive and negative amplitudes, approaching the envelope on both sides. Asymmetry captured deviations from this balance by reducing the amplitude on one side relative to the other, as illustrated in L. Right: tail postures reconstructed from these xy-dynamics. **C:** Reconstruction of the full tail movement from the modeled h- and xy-dynamics and the xy-linked component of h. (i) Left: modeled component time courses; right: the corresponding reconstructed tail postures over time. (ii) The same movement plotted as a trajectory in three-dimensional PC space, revealing its cylindrical geometry. **D:** Example movement comparing observed data and model fit. Observed dynamics are shown as solid lines, with tail postures over time in dark red; modeled dynamics are shown as thick semitransparent lines, with reconstructed postures in gray. The modeled movement was generated from the six parameters shown in A and B. Fit quality was quantified as the *R*^2^ between the observed and modeled traces concatenated across x, y, and h. **E:** Distribution of reconstruction performance across all movements. The Vertical line indicates the *R*^2^ = 0.7 threshold. **F:** Fraction of movements reconstructed with *R*^2^ *>* 0.7 across bout types. Bout types comprising more than 1% of movements in the dataset are shown; remaining bout types are shown in Figure S4C-D. Bout types shown are approach swim (AS), slow1 (S1), long capture swim (LCS), J-turn (JT), high angle turn (HAT), routine turn (RT) and shadow avoidance turn (SAT). **G**,**J:** Effects of the h-dynamic parameters on tail dynamics. In each panel, one parameter is varied across three increasing values. For each value, the corresponding h-dynamics and reconstructed movement are shown. The h-dynamics traces are shown in progressively lighter shades of green, corresponding to increasing parameter values. (G) Max tail bend (measured in a.u.) corresponded to the peak of the h-dynamics (green dot) and determined the most extreme unilateral bend in the reconstructed tail posture (green postures). (J) Increasing time to bend (measured in milliseconds) delayed the rise to the peak without changing the maximum bend amplitude. **H**,**I**,**K**,**L:** Effects of the xy-dynamics parameters on tail dynamics. Each parameter is shown at three increasing values. Line shade denotes parameter value, with the direction of increasing values indicated in each panel. The y-dynamics are shown prominently, whereas the similar x-dynamics are shown faintly in the background. Reconstructed movements are shown as sequences of tail postures on the right. (H) Increasing duration (measured in milliseconds) lengthened the oscillatory epoch. (K) Increasing tail-beat frequency (measured in Hz) increased oscillation frequency without substantially altering overall amplitude or duration. (I) Increasing maximum tail curvature (measured in a.u.) symmetrically increased the positive and negative extrema of the dynamics, strengthening the curvature of leftward and rightward postures. Red and blue vertical lines indicate example time points and their corresponding tail postures. (L) Increasing asymmetry (unitless quantity) reduced the amplitude of oscillations on one side of the dynamics relative to the symmetric envelope (purple line), thereby biasing the oscillation toward the opposite side. Red and blue vertical lines mark opposite phases of the oscillation, corresponding to the maximal leftward and rightward-curved reconstructed tail postures shown on the right.

The xy-dynamics were modeled using a transient oscillator inspired by a burst-spiking model ^29^, which reproduced their oscillatory structure (Figure 2B, see Methods). We summarized these dynamics using four interpretable parameters: duration, tail-beat frequency, maximum tail curvature, and asymmetry. Duration and tail-beat frequency specified the temporal structure of the oscillations, maximum tail curvature specified their overall amplitude, and asymmetry captured differences in oscillation amplitude between the two sides of the tail. The full fitting model included two additional nuisance parameters specifying the timing and phase of the oscillations relative to the detected movement onset (Figure S3B-E, see Methods). Since these two parameters depend on the detected movement onset rather than describing the movement dynamics themselves, we consider the xy-dynamics to be captured by four main parameters. Combining the modeled h- and xy-dynamics together with the modeled xy-linked component of h, the model recapitulated both the empirical tail posture dynamics throughout the movement and the cylindrical trajectories in the posture space (Figure 2C). Although the model treated the h- and xy-dynamics separately, we observed partial coupling during movements with large tail bends. We therefore incorporated this coupling into the model for such movements without introducing additional parameters (18% of the movements, Figure S4A,B; see Methods). Together, the two h-dynamics parameters and four xy-dynamics parameters provided six interpretable parameters that capture the dominant variations underlying hunting movements.

We next asked whether this compact six-parameter model could account for the xy- and h-dynamics of movements in our dataset. For each movement, we fit the model by minimizing the error between the empirical and modeled h- and xy-dynamics (see Methods). The fitted model accurately reproduced both dynamics and the corresponding tail movement over time (Figure 2D). Across the full dataset, 93% of movements were reconstructed with *R*^2^ *>* 0.7 (Figure 2E), indicating that the temporal evolution of most hunting movements can be described within a small interpretable parameter space. To assess whether the model captured movements spanning the full zebrafish movement repertoire, rather than a restricted subset of movement types, we also classified each movement into previously defined movement types using Megabouts ^5,30^ (see Methods). High reconstruction performance was maintained across the dominant bout types observed during hunting (Figure 2F, Figure S4C,D), showing that a broad range of movements can be generated within a shared dynamical framework.

To relate each parameter directly to an observable movement feature, we generated synthetic h, x, and y time series, while varying one parameter at a time and holding the other five fixed. We then reconstructed the corresponding tail movements and examined how each parameter altered specific movement features. In the h-dynamics, increasing peak amplitude increased the maximal unilateral tail bend (max tail bend), whereas increasing the time to bend delayed the rise to peak without altering bend amplitude (Figure 2G,J). In the xy-dynamics, increasing duration prolonged the oscillatory epoch, and increasing tail-beat frequency increased oscillation frequency without affecting movement duration (Figure 2H,K). Increasing maximum tail curvature symmetrically increased oscillation amplitude, strengthening both left- and right-curved postures (Figure 2I). By contrast, increasing asymmetry reduced curvature on one side and biased the oscillation toward the other (Figure 2L). Together, these analyses show that hunting movements can be described and generated by tuning of six continuous and interpretable parameters that jointly shape slow bending and fast oscillatory dynamics.

### Tail-dynamics parameters predict movement outcomes

Each movement was described by six interpretable parameters extracted from tail posture dynamics, whereas its movement outcome could be summarized by three variables^19^: distance traveled (*d*), change in heading angle (Δ*θ*_*head*_), and azimuthal direction of displacement (*ϕ*_*az*_) (Figure 3A, see Methods). Consistent with previous work ^19^, movements occupied a structured movement outcome space in which the fish’s position after the movement covaried with its change in heading angle (Figure 3B). We used this outcome space to ask how distinct parameters of tail movement dynamics shape the behavioral consequences of each movement.

**Figure 3:**
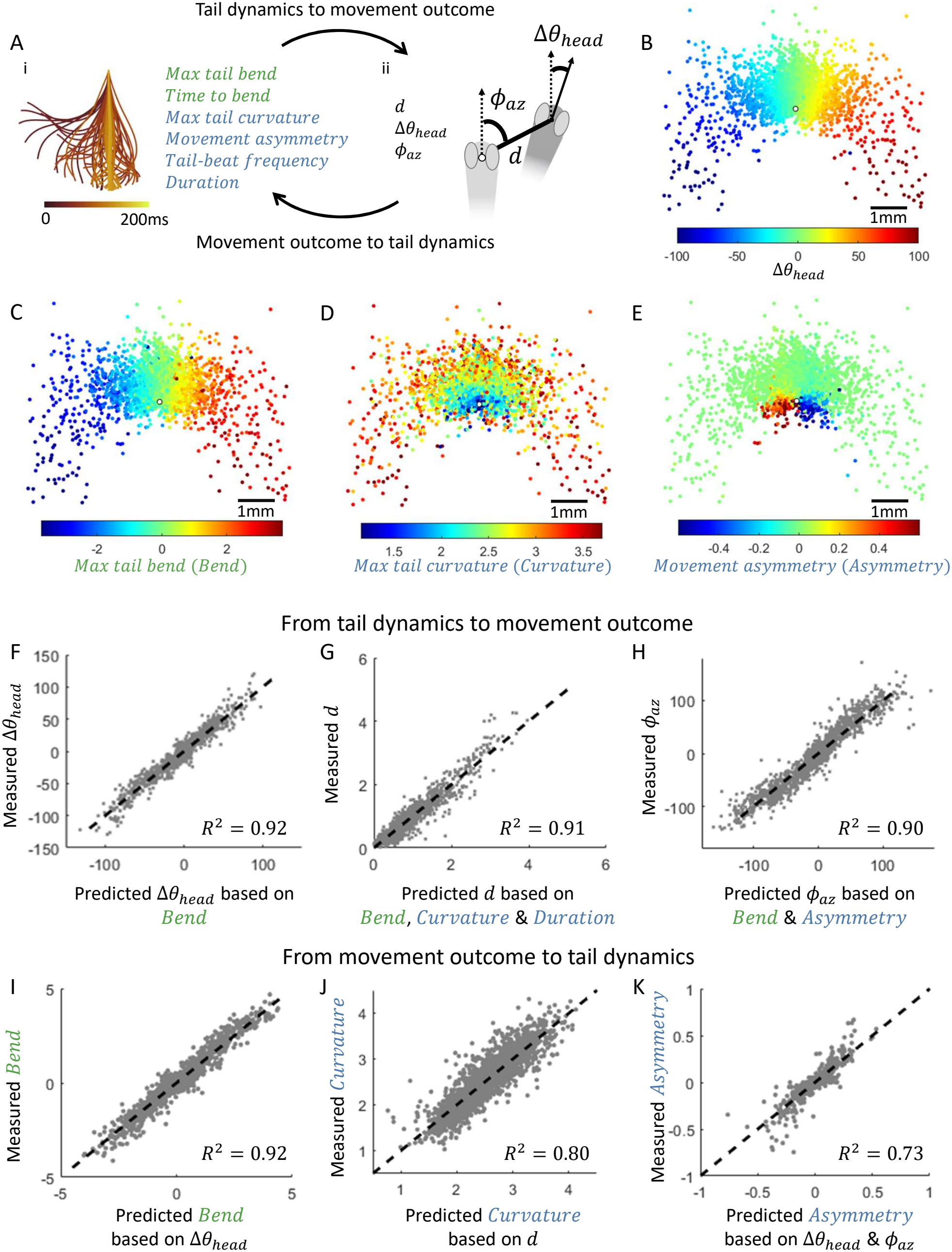
Tail-movement parameters predict and are constrained by movement outcome. **A:** Overview of the six parameters describing tail-movement dynamics and the three variables quantifying movement outcome. Arrows indicate the two directions of analysis: evaluating movement outcome from tail dynamics and estimating tail-dynamics parameters from movement outcome. **B:** Movement outcome repertoire. For each movement, the pre-bout position and heading direction were aligned to a common origin, indicated by the white circle with the heading direction pointing upward. The post-bout position was then plotted relative to this origin as a colored dot. Dots are colored by the change in heading angle, showing that neighboring postbout positions (characterized by *d* and *ϕ*_*az*_) were associated with similar changes in heading angle (Δ*θ*_*head*_). Across the figure, angles are measured in degrees and distance in mm. **C-E:** Movement outcome repertoire colored by individual tail-movement parameters. The white circle indicates the aligned pre-bout position. Outcomes are shown for movements with model fit of *R*^2^ *>* 0.7. (C) Maximum tail bend showed a spatial organization similar to that of change in heading angle. (D) Maximum tail curvature increased with the distance from the pre-bout position. (E) Movement asymmetry was greatest for post-bout positions behind the pre-bout position, suggesting an association between asymmetry and backward displacement. **F-H:** cross-validated prediction of movement outcome from tail-movement parameters. For all analyses in F–K, models were trained on half of the movements from each fish and evaluated on the held-out half, with the procedure repeated after swapping the training and test sets. (F) Predicting the change in heading angle from the maximum tail bend (’Bend’) using a polynomial relationship with powers up to three. (G) Predicting distance traveled based on maximum tail curvature (’Curvature’), movement duration, their interaction, and maximum tail bend. (H) Predicting azimuthal angle from maximum tail bend and movement asymmetry. **I-K:** Cross-validated prediction of tail-movement parameters from movement outcome variables. (I) Maximum tail bend was largely predicted from the change in heading angle, indicating a nearly reciprocal relationship between the h-dynamics and the resulting change in heading angle 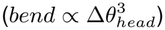. (J) Maximum tail curvature was predicted from distance traveled, indicating that distance provides information about curvature but does not fully determine it (*curvature ∝ log*(*d*)). (K) Movement asymmetry was predicted from the change in heading angle and azimuthal angle (*asymmetry ∝* Δ*θ*_*head*_ + *ϕ*_*az*_). All regressions were performed using linear mixed-effects models with fish identity included as a random intercept to account for inter-fish variability.

This mapping was dominated by a small subset of movement parameters. Maximum tail bend closely recapitulated the organization of change in heading angle across the outcome space (Figure 3C, compare to Figure 3B), predicting changes in heading angle with high accuracy (Figure 3F, see Methods). Adding the remaining five movement parameters only marginally improved predictive performance, whereas these parameters alone had little predictive power for change in heading angle (Figure S5A-D). Together, these results indicate that the change in heading angle was mainly associated with the degree of tail bend.

Maximum tail curvature (’curvature’) showed a clear organization along the distance axis: movements with low curvature produced short displacements, whereas movements with higher curvature produced longer displacements (Figure 3D). We therefore asked which movement parameters predicted variation in distance traveled. Maximum tail curvature, movement duration, and maximum tail bend together predicted distance traveled with high accuracy (*R*^2^ = 0.91) (Figure 3G). Maximum tail bend contributed to the distance largely indirectly, through its association with changes in heading angle in large turns (Figure S5E-F). After accounting for this turn-related component, distance was primarily associated with the maximum tail curvature and duration of the oscillatory tail dynamics, while the remaining model parameters provided little additional predictive power (Figure S5G).

Asymmetry was associated with the azimuthal direction of displacement, particularly for movements covering short distances (Figure 3E). Across all movements, asymmetry together with maximum tail bend predicted azimuthal direction with high accuracy (*R*^2^ = 0.90) (Figure 3H), whereas the contributions of the other parameters were small (Figure S5H,I). Most movements showed symmetric oscillations, for which the azimuthal angle was largely accounted for by maximum tail bend due to the approximately linear relation between the change in heading angle and azimuth (Figure S5H). As oscillation asymmetry increased, the azimuthal angle increasingly deviated from the linear relationship, producing progressively more backward-directed movements. Overall, although six parameters were required to fully describe movement dynamics, movement outcome was largely predicted by four parameters: maximum tail bend, maximum tail curvature, movement duration, and asymmetry. These parameters mapped onto distinct aspects of movement outcome: bend primarily to change in heading angle, curvature and duration to distance traveled, and asymmetry to azimuthal direction.

We next asked whether a given movement outcome is achieved by a unique tail movement or by several possible tail-dynamics patterns. To test this, we inverted the mapping and asked how accurately each movement parameter could be predicted from the outcome variables. Change in heading angle accurately predicted maximum tail bend (Figure 3I), indicating that the degree of tail bend was largely recoverable from the resulting change in heading angle. Distance traveled predicted maximum tail curvature with substantial accuracy (*R*^2^ = 0.80) (Figure 3J), while change in heading angle together with azimuthal direction predicted asymmetry (*R*^2^ = 0.73) (Figure 3K). Movement duration was less accurately predicted (Figure S6A). Although tail-beat frequency did not improve the prediction of movement outcome once the other parameters were included (Figure S5D,E,G,I), its variation could nevertheless be predicted to a substantial degree from the outcome variables and showed strong fish-dependent variation (Figure S6A-C). Thus, movement outcome carried substantial information about multiple tail-dynamics parameters. However, recovery was most complete for maximum tail bend, whereas the oscillatory xy-dynamics parameters were constrained, but not uniquely determined, by the measured outcome.

### The hunting repertoire forms a constrained continuum of interpretable dynamics

Having established that a compact set of parameters can reconstruct individual hunting movements and explain their outcomes, we next asked whether hunting movements occupy a constrained region of tail-dynamics space, and whether gradual changes within this space correspond to interpretable changes in the underlying movement dynamics. We first examined the relation between the parameters of the xy- and h-dynamics. Rather than uniformly sampling this space, movements formed a bounded repertoire in which some parameter combinations were common while others were rarely observed (Figure 4A-C, Figure S7A-K). To provide a familiar reference for interpreting this space, we colored movements according to their conventional bout-type assignments ^30^ (see Methods). Bout-type identity largely tracked variation in maximum tail bend. To connect this parameter-space organization to the underlying movements, we examined representative movements along the maximum tail bend and duration axes (Figure 4D). Nearby movements showed similar tail movement and posture dynamics, whereas gradual shifts across these axes corresponded to graded changes in bend magnitude, bend direction, and movement duration.

**Figure 4:**
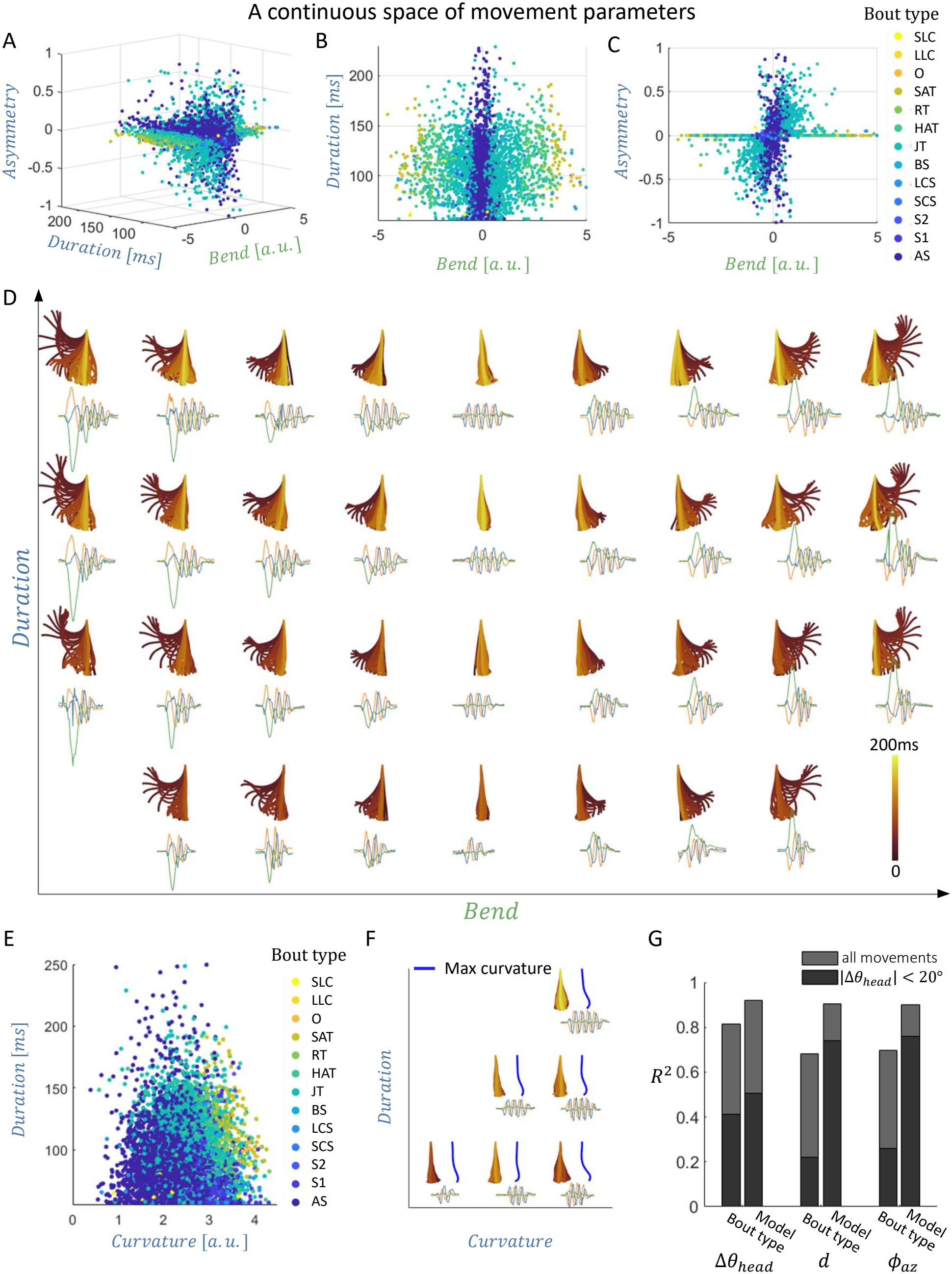
The hunting movement repertoire forms a constrained continuum in tail-dynamics parameter space. **A-C:** Three-dimensional space of movements with model fit *R*^2^ *>* 0.7 defined by maximum tail bend, movement duration, and movement asymmetry. Movements are colored and ordered by cluster type ^30^, showing that bout types were distributed gradually along the maximum tail bend axis. **D:** Example tail movements and dynamics positioned along the maximum tail bend and duration axes, illustrating how variation in the model parameters corresponds to interpretable changes in tail movement. Position along the maximum tail bend axis ranged from strong leftward to strong rightward bending, whereas increasing duration produced progressively longer movements. **E:** Movements positioned along the duration and maximum tail curvature axes and colored by movement type. These parameters were partially dependent, such that movement duration constrained the range of maximum tail curvature. **F:** Example tail movements and dynamics of movements with low bend values positioned along the maximum tail curvature and duration axes, illustrating graded changes in tail movement. Moving along the maximum tail curvature axis increased the most curved posture during the movement, shown in blue, whereas increasing duration produced progressively longer movements **G:** Prediction of movement outcome based on either the model parameters or conventional bout types. Both approaches predicted changes in heading angle with similar accuracy, for small turns and across all movements. Model parameters provided more accurate predictions of distance and azimuth angle than bout type, particularly for small turns. Bout types shown are approach swim (AS), slow1 (S1), slow2 (S2), burst swim (BS), short capture swim (SCS), long capture swim (LCS), J-turn (JT), high angle turn (HAT), routine turn (RT), shadow avoidance turn (SAT), O-bend (O), long-latency C-start (LLC) and short-latency C-start (SLC).

We next examined whether this constrained, continuous organization was also evident within the xy-dynamics. The relationship between xy-dynamics parameters, such as maximum tail curvature and duration, also formed a bounded structure (Figure 4E) with gradual shifts across these axes corresponding to continuous changes in movement dynamics (Figure 4F). Bout-type identity also shifted gradually across this space, although neighboring bout types overlapped more extensively than along the maximum tail bend axis. Mapping the relation between bout-type identity and the model parameters showed that cluster identity mostly explained variance in maximum tail bend and maximum tail curvature (Figure S7L). The continuous model parameters predicted movement outcomes more accurately than bout-type identity, particularly for small turns (Figure 4G). Together, these results show that the hunting repertoire is organized by continuous but constrained combinations of co-occurring tail-dynamics parameters.

### The tail-dynamics model generalizes across age groups and key parameters can be extracted directly from kinematics

Having identified the tail-dynamics parameters that organize the repertoire and explain movement outcome, we first asked whether the model generalized to an independent dataset of hunting movements from 13-15 dpf fish (25 fish, 5,006 movements). Despite an increase of up to 30% in body length (see Methods) and developmental changes in hunting behavior ^31^, movement reconstruction performance was similar to that observed in 5-7 dpf fish (Figure 5A). Moreover, the relationships between movement parameters and outcome were preserved: the fitted parameters explained a similar proportion of variance in movement outcomes, and the corresponding parameters could likewise be estimated from movement outcomes with comparable accuracy (Figure S8A,B, see Methods). Thus, both the model and the mapping between movement dynamics and outcome generalized across developmental stages.

**Figure 5:**
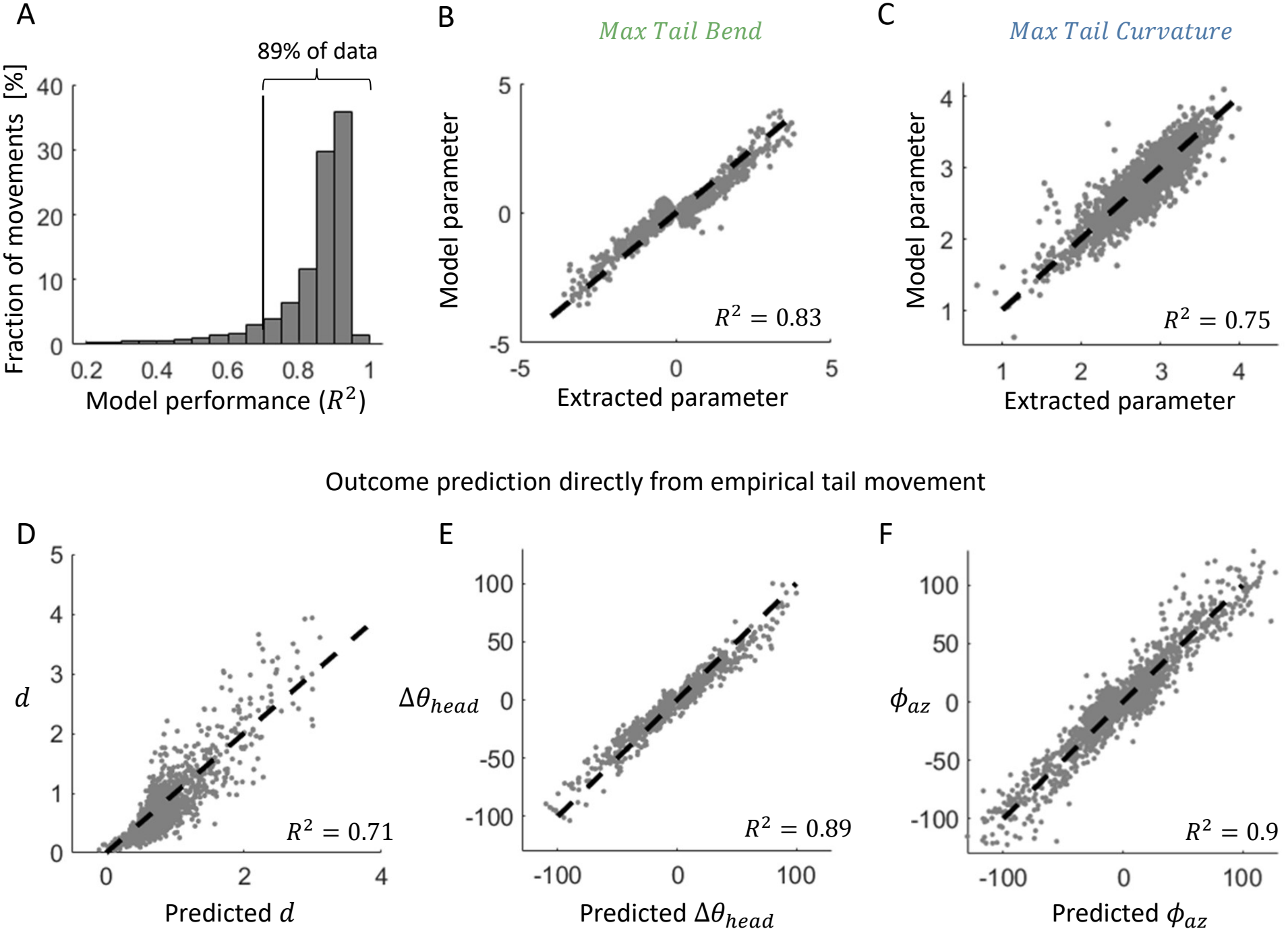
Tail dynamics generalize across age groups and key parameters are directly measurable from kinematics. **A:** Distribution of reconstruction performance across all movements in the 13-15 dpf dataset. Reconstruction performance was comparable to that observed in 7 dpf fish (compare with Figure 2E), supporting the applicability of the same dynamical model across age groups. **B:** Maximum tail bend extracted directly from empirical tail kinematics in the 13–15 dpf dataset, without fitting the dynamical model, compared with the maximum tail bend parameter calculated by the model, showing reliable extraction across fish. The close correspondence between the two measures indicates that maximum tail bend can be reliably estimated directly from tail kinematics. **C:** Maximum tail curvature extracted directly from empirical tail kinematics, in the same 13-15 dpf dataset, without fitting the dynamical model, compared with the fitted maximum tail curvature model parameter. The two measures were strongly related, indicating that maximum tail curvature can be estimated directly from tail kinematics. **D-F:** Prediction of individual outcome parameters from empirically extracted estimates of the model parameters, showing that movement outcomes can be accurately predicted without fitting the dynamical model.

Because the key model parameters corresponded to specific, interpretable features of the tail dynamics (Figure 2G-L), we next asked whether they could be estimated directly from tail kinematics without fitting the model. Leveraging the robustness of posture PCs and rotation across fish (Figure S1), we used this shared posture representation to decompose empirical movements into h- and xy-dynamics and derived empirical estimates of maximum tail bend, maximum tail curvature, and asymmetry. We evaluated the relationship between these empirical measures and the corresponding fitted model parameters using the 5-7 dpf dataset, and independently validated the resulting mappings in the 13-15 dpf dataset (see Methods). Maximum tail bend and maximum tail curvature showed high agreement with the fitted values (Figure 5B,C), whereas asymmetry showed more moderate agreement (Figure S8C). Nevertheless, the empirically extracted parameters accurately predicted movement outcomes (Figure 5D-F). Thus, the interpretability of the model allowed us to translate its fitted parameters into practical measurements that can be extracted directly from behavior and used to estimate movement outcomes. More broadly, these measurements provide interpretable behavioral variables for future analyses of the neural control of movement.

## Discussion

Natural movement repertoires can appear complex because individual movements vary across multiple kinematic features. By resolving larval zebrafish hunting movements into their underlying dynamics, we found that this diversity arises from the simultaneous expression of two components: a slow, non-oscillatory bending dynamic and a fast, oscillatory undulatory dynamic. A compact model described these dynamics using six continuous, interpretable parameters, accurately reconstructed movements across the repertoire, and generalized to an older developmental stage. Specific parameters were associated with distinct movement outcomes: tail bend with changes in heading angle, oscillation curvature and duration with displacement distance, and oscillation asymmetry with azimuthal displacement direction. Conversely, movement outcome strongly constrained, but did not always uniquely specify, the underlying tail dynamics. Hunting movements formed a bounded continuum within an interpretable dynamical space. Finally, we provide a practical approach to extract key parameters directly from empirical tail movements, providing a low-dimensional, interpretable description of individual movements. These parameters also offer candidate behavioral variables for linking natural movement dynamics to their neural control.

Our work uncovered two co-occurring tail dynamics: slow, non-oscillatory bending dynamics and fast oscillatory dynamics. For most movements (82%), these two dynamics could be described as independent. However, in movements with very large tail bends (18% of movements), model fits improved when oscillatory frequency was transiently coupled to the bending timescale (Figure S4A,B). A second form of dependence was apparent across the full repertoire in the relationship between oscillatory and bending amplitudes: as maximum tail bend increased, the range of observed oscillatory curvatures became more restricted (Figure S7G). Together, these observations suggest that the two dynamics may be generated largely independently, while their simultaneous expression is constrained, potentially by neural or biomechanical factors, with transient coupling emerging specifically during large tail bends.

Known motor circuits provide plausible substrates for both the separation and transient coordination of these dynamics. Distinct descending commands engage different spinal motor pools: excitatory V2a neurons mediating rapid steering excite ultra-fast primary motoneurons, whereas V2a neurons mediating slow exploratory movements excite secondary, but not primary, motoneurons ^32^. Consistent with this functional separation, ventromedial spinal projection neurons introduce asymmetric tail deflections into the underlying swimming pattern ^33^, whereas MLR-recruited V2a reticulospinal neurons regulate forward-swimming kinematics ^34^. Individual neurons in the nucleus of the medial longitudinal fasciculus (nMLF) are also preferentially linked to bout duration and tail-beat frequency during forward swimming ^35^. Population recordings further revealed a low-dimensional reticulospinal organization in which speed- and steering-related modules act combinatorially across locomotor space^36^. The ventromedial turning pathway may also provide a mechanism for transient coordination: its ablation reduced the large initial tail deflection while sparing subsequent symmetric undulations and shortened the prolonged first oscillation cycle of turns, suggesting that bend-related commands can transiently slow the oscillatory rhythm ^33^. Whether the relationships between the two dynamics reflect coordinated neural control, biomechanical constraints, or both, remains unresolved.

We characterized the slow bending dynamics with two parameters: maximum tail bend and time to bend. Of these, the maximum tail bend was tightly linked to turning outcome: it accurately predicted the change in heading angle and could, in turn, be recovered from the change in heading angle with similar accuracy (Figure 3F, I, Figure S8A). This close bidirectional relationship suggests that maximum tail bend is a principal kinematic variable associated with change in heading angle, largely independently of the other movement parameters. This relationship is consistent with a steering command whose magnitude is reflected in both bend amplitude and the resulting turn. Consistent with this interpretation, steering-related neural activity scales with turn magnitude, and activation of descending steering pathways can produce graded ipsilateral tail deflections ^33,37^. Directional swims also shift a persistent representation of heading in the anterior hindbrain by an amount corresponding to the executed turn, likely through motor-efference signals^38^. Our results suggest that maximum tail bend may provide a compact kinematic variable linking the specification and execution of a turn to the subsequent updating of internal heading.

The fast oscillatory dynamics were characterized by four parameters: maximum tail curvature, tail-beat frequency, duration, and asymmetry. Curvature and duration jointly predicted movement distance (Figure 3G, S5E-G). However, the inverse mapping was weaker: movement distance recovered most of the variance in maximum tail curvature (Figure 3J) but only moderately recovered duration (Figure S6A). This difference between the forward and inverse mappings indicates that achieving a particular displacement does not require a unique set of oscillatory parameters. Rather, movement distance strongly constrains maximum tail curvature while leaving residual variability in both curvature and duration, such that similar displacements can be produced by multiple curvature-duration combinations. This flexibility is nevertheless constrained: a given maximum tail curvature was associated with only a restricted range of possible durations (Figure 4E). Thus, distance constrains the oscillatory dynamics to a bounded set of curvature-duration combinations, allowing similar displacements to be achieved over different timescales without specifying a unique movement.

While outcome could be predicted from four model parameters, tail-beat frequency did not explain additional variance (Figure S5E,G). Nevertheless, frequency itself could be predicted from outcome (Figure S6B). Thus, frequency covaried systematically with movement outcome but carried little unique information about distance once curvature and duration were taken into account. Moreover, the relationship between frequency and outcome was strongly fish-dependent, suggesting that frequency may also be influenced by factors beyond movement outcome. Consistent with this possibility, tail-beat frequency and movement duration were shown to vary with environmental temperature ^39^, indicating that temporal features of swimming can be adjusted according to external conditions. Whether such modulation is restricted primarily to temporal parameters or also extends to parameters more tightly associated with movement outcome, such as maximum curvature, remains unclear. Together, these observations raise the possibility that movement outcome and its detailed execution are partly separable, which may allow similar displacements to be produced through different combinations of oscillatory parameters across individuals or environmental conditions.

The dynamical description also raises the question of when movement parameters are specified to generate a movement. Curvature, frequency, and asymmetry can be estimated from the initial undulation, and the bending dynamics can likewise be characterized from their early evolution, indicating that much of the structure distinguishing movements is already apparent early in their execution. This is consistent with previous observations that turning and forward swimming differ primarily during the first tail-undulation cycle, whereas subsequent undulations are nearly identical ^33^. Together, these observations raise the possibility that key movement parameters are specified by an early motor command rather than controlled during movement execution. Such a strategy is consistent with the impulse-like input used in our model (Figure S3B), although other forms of motor command preceding the movement are also possible. Future neural measurements could test whether premotor activity specifies these movement parameters before their expression in the evolving tail dynamics.

A continuous repertoire can coexist with recognizable movement types, with categorical and continuous descriptions capturing complementary aspects of its organization. In our dataset, bout types occupied distinct but over-lapping regions of the continuous tail-dynamics space, with cluster identity reflecting variation most strongly in maximum tail bend and, to a lesser extent, in other movement parameters. The dynamical description captures variation within movement types and graded transitions between them, provides a direct account of how individual movements differ, and enables more accurate prediction of each movement’s displacement and change in heading angle than bout identity alone. This may be particularly useful when comparing development, context, or perturbations, because two conditions could show similar usage of a given bout type while differing systematically in its bend, curvature, duration, or asymmetry ^39^. By preserving this graded variation in interpretable movement variables, the dynamical representation may also facilitate identifying neural signals that modulate particular aspects of movement, which could be obscured by categorical bout labels.

An additional benefit of the dynamical framework is that it provides a principled basis for resolving the temporal organization of prolonged tail movements. Successive movements can occur with little or no inter-bout interval, making it difficult to distinguish a single prolonged movement from several movements concatenated into a compound movement. Such compound movements have been reported previously and become more frequent over development ^40^. In our framework, each movement has an oscillatory envelope that rises at movement onset and decays toward its end. This expected temporal structure provides a way to identify the constituent components of compound movements: a new movement initiated before the preceding envelope has closed may be revealed by a renewed expansion of the envelope, or with an abrupt change in bend or oscillatory structure. Consistent with this interpretation, we observed compound movements whose components were separated by only a brief interval or concatenated without an intervening pause (2.5% of movements, Figure S8G-I). These included repeated large-bend movements, successive forward movements, and combinations of forward and turning movements. Thus, repertoire diversity arises not only from continuous variation in the dynamics of individual movements, but also from the sequential composition of individual movement components into compound movements. Future work could use this framework to automatically identify such movement components, characterize each component in terms of the model parameters, and determine how patterns of movement concatenation change over development.

Despite substantial changes in hunting behavior over development ^31^, the same dynamical description generalized to an older developmental stage. The preservation of the dynamical description and parameter-outcome relationships suggests that the basic coordinates of the hunting repertoire may remain stable across development. Developmental changes in behavior could therefore occur within this shared dynamical space, rather than requiring a different set of underlying movement dynamics. Determining whether this framework remains stable across a broader developmental range will require extending the analysis to additional developmental stages.

Whether this organization is preserved across different experimental conditions or other fish species^41^ remains to be established. Extending the framework to head-fixed preparations could offer two complementary approaches to quantify the behavior. First, extracting interpretable tail-dynamics parameters could facilitate identifying neural signals involved in controlling specific aspects of movement. Second, using these parameters to estimate the displacement and changes in heading angle that tail movements would produce during free swimming could provide more informative behavioral readouts and enable more precise closed-loop experiments. More generally, although tail dynamics accurately predicted movement outcomes in our dataset, additional motor components, such as pectoral fins ^42,43^, may also contribute to movement outcome. Incorporating these components could reveal whether they explain additional variation in displacement and orientation beyond that captured by tail dynamics alone.

The dynamical system used here should not be interpreted as a unique mechanistic account of tail-movement generation. Other formulations capable of producing transient bending and decaying oscillations could potentially reconstruct the same movements. These formulations may imply different mechanisms for generating these dynamics and may also identify somewhat different parameters for describing them; for example, one formulation might represent oscillatory amplitude as maximum tail curvature, while another might use maximum tail sweep (Figure S6D-F). Distinguishing among such mechanistic descriptions will ultimately require neural measurements that constrain how these dynamics are generated.

Despite this non-uniqueness, the model provided two main contributions. First, it identified interpretable dynamical parameters that could subsequently be extracted directly from empirical tail movements without fitting the generative model, while preserving their relationships with displacement and changes in heading. Second, the model demonstrated that six parameters are sufficient to reconstruct the temporal evolution of the two dynamics across the repertoire. While tail-beat frequency and movement duration are commonly used, reduced versions of our model revealed distinct contributions for maximum tail bend, asymmetry, and maximum tail curvature to movement reconstruction (percentage of movements with *R*^2^ *>* 0.7: 67%, 86%, and 90% when fixing each parameter, respectively). Importantly, these parameters were also strongly predictive of movement outcomes, indicating that they capture behaviorally meaningful variation beyond what is reflected in reconstruction accuracy.

Together, these findings show that the diversity of the zebrafish hunting repertoire emerges from continuous but bounded combinations of shared bending and oscillatory dynamics. Linking these dynamics to behavioral outcome and to experimentally measurable parameters provides a bridge from repertoire-level organization to testable questions about motor-circuit control.

## Methods

### Experimental procedures

#### Zebrafish maintenance

Nacre zebrafish (*Danio rerio*) embryos expressing elavl3:H2B-GCaMP6s^44^ were collected according to established procedures ^45^ and raised at 27°;C with a light and dark cycle of 14/10 h. Larvae were fed live rotifers (Brachionus plicatilis) daily from 5 dpf. All procedures were performed with approval by the Hebrew University of Jerusalem Animal Ethics Committee.

#### Imaging natural hunting behavior

Individual larvae, 5-7 dpf (n=30) and 13-15 dpf (n=25), were placed in a 20-mm-diameter, 2.5-mm-deep transparent-bottom chamber (CoverWell Imaging Chambers, catalog no. 635031, Grace Bio-Labs) containing 30 *Paramecia* and allowed to hunt freely for 15 min. The chamber was illuminated from above with white visible light, while infrared LEDs (850 nm; LDR2-100IR2-850-LA, powered by PD3-3024-3-PI; CCS Inc., Kyoto, Japan) were positioned beneath the chamber. Room temperature was maintained at 27°;C. Larval behavior was recorded from above using a high-speed camera (Mikrotron 4CXP) at 500 frames/s with a spatial resolution of 45 *µm* per pixel.

### Fish tracking

Recordings of natural hunting behavior were analyzed post hoc using a custom algorithm based on computervision and graph-theory methods. The algorithm detected the fish, extracted its morphological features and identified hunting events and movement bouts.

#### Fish detection

To detect the fish and extract its features, we segmented the fish from its surroundings on a frame-by-frame basis while excluding prey items and other objects, such as dust particles. The imaging-chamber contour was first identified based on its large, elliptical convex hull and removed. Following removal of the chamber contour, the fish constituted the largest object in the frame. Its identity was further verified based on its overall shape and size.

#### Fish feature extraction

Fish features were extracted as previously described ^19^, with the principal steps summarized below. Once the fish was identified, we computed its contour and searched for objects contained within it. The eyes were identified as the two darkest objects within the fish contour that were elliptical, adjacent to one another, and within a predefined size range. An ellipse was fitted to each eye contour, providing its center and orientation, defined by its major and minor axes. Fish heading was defined by a vector originating at the midpoint between the two eye centers, orthogonal to the axis connecting them, directed away from the tail. The angle between the fish heading and the major axis of each fitted eye ellipse was used to identify eye convergence and divergence. The tail midline was extracted by applying morphological transformations to the fish contour until it was reduced to a single-pixel-wide line. The resulting skeleton was represented as an ordered sequence of points extending from the swimbladder to the tail tip. The swimbladder was identified based on the contrast along the midline, exploiting the fact that in our setup the swimbladder appeared as a bright circle surrounded by a darker circumference. We therefore searched along the midline for two darker points at the expected position of the swimbladder, based on the median tail-tip-to-swimbladder distance estimated separately for each fish. Tail-tip-to-swimbladder distance was 3.33 *±* 0.16mm across the 5-7 dpf fish and 3.74 *±* 0.29mm for the 13-15 dpf fish.

#### Hunting-event and bout detection

Hunting events and movement bouts were identified using the procedures described previously ^19^. Briefly, hunting events were identified from periods of eye convergence, and individual movement bouts were detected from tail-tip velocity. The same detection criteria and thresholds described in ^19^ were used here.

### Extracting tail-posture dynamics

#### Posture description

To obtain a fixed-length representation of tail posture, we first converted the tracked tail in each frame into a standardized set of points. Because the number of pixels composing the tracked tail varied with its posture, we standardized tail length by fitting the x- and y-coordinates separately as seventh-degree polynomial functions of position along the tail and sampling the fitted curves at 105 equally spaced positions. Interpolation was less reliable near the tail tip, so the final six points were excluded, leaving 98 segments between the remaining points. We then calculated the angle of each segment relative to the horizontal axis of the image. To describe tail shape independently of the fish’s overall orientation, in each frame we subtracted the mean segment angle from all segment angles. This removed the common rotational component while preserving the relative angles between segments that define tail posture as a 98-angle posture vector for each frame. Based on this vector, tail curvature (Figure 1O) was calculated per frame as Σ_*i*_(|*α*_*i*+1_ *− α*_*i*_|) where *α*_*i*_ is the angle of segment *i*, and tail tip angle (Figure 1L) was calculated as *α*_98_ *− α*_1_.

#### Low-dimensional description of tail postures

We concatenated the resulting angles across frames into a frames *×* segment-angle matrix *A* and performed singular value decomposition (SVD) on frames associated with detected movements. For each detected movement, the analysis window was extended by 30 frames before and after the movement to allow the dynamics to decay toward a near-zero baseline for subsequent spectral analyses. For each fish, we concatenated the tail-segment angles from these extended movement windows into a single frames *×* segment-angle matrix *A* and performed singular value decomposition (SVD). We decomposed the matrix as *A* = *U* Σ*V* ^*T*^, where the columns of *V* describe posture components across tail segments and the corresponding columns of *U* describe their temporal coefficients. The first three posture components captured the dominant variation in tail posture and were retained for subsequent analyses (Figure 1D,G, Figure S1A). Thus, each tail posture was represented by three corresponding coefficients defining its position within the three-dimensional posture space. The three posture components showed a consistent ordering across fish, with components of similar shape appearing in the same rank order across individuals. To establish a consistent sign convention across fish, each principal posture component and its corresponding temporal coefficients were flipped, when necessary, such that the value of the first tail segment was positive (Figure S1B).

#### Transformation to cylinder-aligned coordinates

We next transformed the three-dimensional coordinates describing each posture into a coordinate system aligned with the cylindrical organization of the movement repertoire, with x and y spanning the cylinder’s cross-sectional plane and h following its longitudinal axis (Figure 1F, Figure S1E-H, J-L). We therefore sought three directions in the three-dimensional principal-component space that separated the temporal dynamics of tail posture coefficients (Figure 1Eii).

To identify these directions, we optimized a 3 *×* 3 linear transformation of the three principal-component coefficient time series (Figure 1Eii) to minimize the sum of squared pairwise cross-correlations between the resulting time courses. The optimization process was performed over a 3 *×* 3 matrices composed of unit-norm vectors using Matlab’s *fminsearch* function. This procedure identified two directions with distinct temporal dynamics: one direction followed the longitudinal axis of the cylinder and was defined as *h*, whereas a second direction lay within the cylinder’s cross-sectional plane and was defined as *x*. These two directions were not constrained to be orthogonal (Movie 1).

We then identified the remaining direction within the cross-sectional plane. For this purpose, we used movements with small changes in heading angle (Δ*θ*_*head*_ *<* 5°;), which showed minimal displacement along the longitudinal direction, to estimate the cylinder’s cross-sectional plane. For each movement, we focused on the posture-space points lying along the circumference of the cylinder and calculated a vector normal to the plane defined by these points. The cross-sectional plane was then estimated by averaging these normal vectors across movements. Importantly, this average vector was not parallel to the h-axis direction, meaning that the xy-plane was not per-pendicular to the h-dynamics. The *y* direction was defined to be orthogonal to this average vector and to the x-direction, ensuring that it lay within the cross-sectional plane. Finally, the x- and y-axes were rotated together within the cross-sectional plane by a fixed angle selected in the representative fish to align the y-axis with the direction along which the amplitude truncation associated with coupled movements was maximal (Figure S4A-B, see Coupled dynamics in Methods section). The complete transformation was calculated for the representative fish and then applied unchanged to all fish in the dataset, yielding the *x*-, *y*-, and *h*-dynamics used throughout the subsequent analyses. The resulting transformation matrix was:

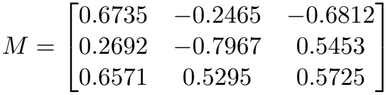

such that the cylinder-aligned coefficients over time (Figure 1H) were given by *MU* ^*T*^, where *M* is the transformation matrix defined above.

#### Scaling posture dynamics across fish

Because the temporal SVD components (the columns of the U matrix) are normalized to unit length, the scale of values in each timestep depends on the number of frames included in the decomposition, which in turn depends on the number and duration of movements. The cylinder-aligned dynamics derived from these components were therefore not directly comparable in amplitude across fish. To compensate for this normalization and enable comparison across fish, we multiplied the x-, y-, and h-dynamics of each fish by that fish’s second singular value. The same scaling factor was applied to all three dynamics, preserving similar amplitudes across them. We used the second rather than the first singular value because the first posture component was dominated by slower, turn-related dynamics, and its singular value could therefore vary with the prevalence and magnitude of turning movements across fish, rather than reflect an overall scale of posture dynamics (Figure S1D).

#### Removing the xy-linked component from h-dynamics

Although the cylinder-aligned transformation largely separated the slow h-dynamics from the faster xy-dynamics, a small high-frequency residual remained in the hdynamics. The phase of this residual was systematically related to the phases of the x- and y-dynamics (Figure S2A), suggesting that it reflected a small xy-linked contribution. We therefore estimated, separately for each fish, the effect of the x- and y-dynamics on the h-dynamics and subtracted it from the h-dynamics before fitting the h-dynamics with the dynamical model.

To estimate the xy-linked contribution, we focused on movements in which the slow bending dynamics were expected to be near zero: movements with an absolute change in heading angle smaller than 5°;. For these movements, we assumed that the dynamics of *h*(*t*) were dominated by the xy-linked contribution and could therefore be predicted by the x- and y-dynamics. The x-, y-, and h-dynamics of these movements were transformed into Fourier space. We then estimated fish-specific frequency-domain filters, 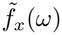 and 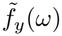, that predicted the measured h-dynamics from the corresponding x- and y-dynamics (Figure S2B). All movements were padded to 256 samples in time, such that both 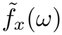 and 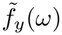 were 256-sample filters estimated separately for each fish. At each frequency, we applied a robust regression across movements to estimate the contributions of x and y according to:

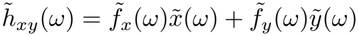

using Python’s HuberRegressor with default parameters. Once fitted, we used the same 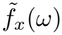 and 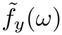 to calculate 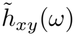 for all movements. The resulting estimate of the xy-linked component, 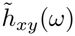, was transformed back into the time domain to obtain *h*_*xy*_(*t*). For each movement, the cleaned h-dynamics were then calculated as

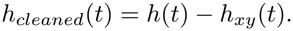

Removing the xy-linked component yielded cleaned h-dynamics dominated by a slow excursion to a peak or trough followed by a return toward baseline (Figure S2C-D), while reducing both their high-frequency content (*>* 20 Hz) and their correlations with the x- and y-dynamics (Figure S2E-G). The relative contribution of the removed component was greatest for movements with weak h-dynamics and decreased as the magnitude of the h-dynamics increased (Figure S2H), indicating that the correction primarily removed a small oscillatory residual rather than the dominant slow h-dynamics. The resulting *h*_*cleaned*_ trajectory was used for all subsequent fitting of the h-dynamics.

### Modeling h-dynamics

#### Model dynamics and parameters

After removal of the xy-linked component from the h-dynamics, the cleaned h-dynamics were characterized by a single excursion to an extermum followed by a return to baseline (Figure S2C-D; Figure S3A). We modeled the initial excursion using a cosine function, followed by an exponential decay to baseline:

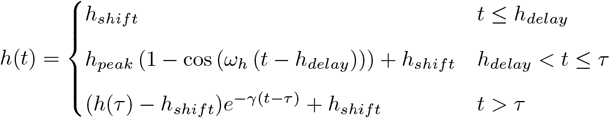

where

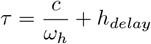

The two main parameters controlling the shape of the h-dynamics were *h*_*peak*_ and *ω*_*h*_ (Figure S3A). The parameter *h*_*peak*_ set the magnitude and direction of the h-dynamics, with the signed peak excursion of the cosine component equal to 2*h*_*peak*_. The parameter *ω*_*h*_ controlled their temporal scale, with larger *ω*_*h*_ producing a faster rise and earlier peak. Equivalently, 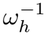 provided a measure of the characteristic timescale of the bending dynamics.

The model additionally contained *h*_*delay*_, which specified the timing of the modeled h-dynamics relative to the detected movement onset, and *h*_*shift*_, which accounted for a small initial offset from baseline posture. These latter two parameters described the alignment and initial state of the modeled dynamics rather than the movement dynamics themselves.

The transition from the cosine segment to the exponential decay occurred at a fixed cosine phase, determined by *c*. The direction of the h-dynamics was determined from the sign of the integral of the observed h-dynamics over the movement, and *c* was assigned accordingly (*c* = 1.624*π* for a positive integral and *c* = 1.465*π* for a negative integral). The exponential decay rate was fixed at *γ* = 45*s*^*−*1^ for all movements.

#### Fitting the h-dynamics

The four free parameters of the h-model, *h*_*peak*_, *ω*_*h*_, *h*_*delay*_, and *h*_*shift*_, were fitted using SciPy’s *curve fit* function. Initial values for *h*_*delay*_ and *ω*_*h*_ were 1 ms and 125 rad/s, respectively. The initial value of *h*_*shift*_ was defined as the mean of the first and last values of the h-dynamics. *h*_*peak*_ was initialized as half the larger absolute deviation of the observed h-dynamics from this baseline, retaining the sign of that deviation. *ω*_*h*_ was constrained to values greater than 50 rad/s. Each movement was fitted twice, constraining *h*_*shift*_ to be positive or negative, respectively, and the solution with the higher explained variance was retained. When the sign of the initial guess for *h*_*shift*_ was incompatible with the corresponding constraint, the initial guess was set to zero.

### Modeling xy-dynamics

The xy-dynamics consisted of transient oscillations whose duration, frequency, amplitude, and asymmetry varied across movements. We therefore constructed the xy-model in four stages (Figure S3B): (i) generation of the oscillatory dynamics, (ii) addition of asymmetry, (iii) adjustment of oscillation amplitude, and (iv) addition of a decay to baseline.

#### Generating the oscillatory dynamics

The xy-dynamics were modeled using a transient oscillator inspired by the canonical “2+1” Izhikevich model of bursting neurons ^29^:

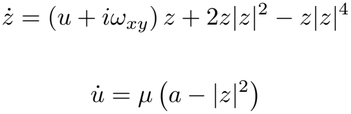

Here, *ω*_*xy*_ denotes angular frequency in rad/s; the corresponding dynamics frequency in Hz is given by *f*_*xy*_ = *ω*_*xy*_*/*2*π*.

Because the original Izhikevich system generates recurrent transient oscillations autonomously, it does not provide a defined trigger for the onset of an individual oscillatory episode (a movement). We therefore introduced a delta-function input to the u-dynamics to initiate each modeled movement and held u fixed before this input. The delta-function input was implemented numerically by setting *u* = *u*_*str*_ at *u*_*time*_. Therefore, the u-dynamics evolved according to the model equations:

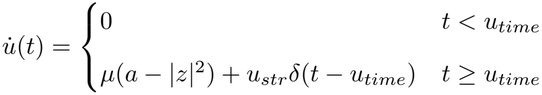

The complex variable *z* generated the oscillatory dynamics, with

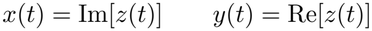

The two main parameters at this stage were *ω*_*xy*_, which controlled the oscillatory frequency of the xy-dynamics, and *u*_*str*_, which controlled the duration of the transient oscillations through the u-dynamics (Figure S3B,D). Two additional parameters specified the timing and initial phase of the oscillation: *u*_*time*_, the timing of the input relative to the time *t* = 0 of the movement data (defined as 30 frames before the detected movement onset), and *z*_0_, the initial phase of *z*. Importantly, there was a delay between the time of the input (captured by *u*_*time*_) and the time of movement onset, where the xy-envelope (defined by 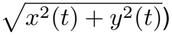 increased sharply from its near-zero values. Lastly, we assumed *u*(*t*) = 0 before the delta-function input and initialized

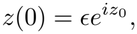

with *ϵ* = 10^*−*7^. The constants *µ* = 0.3 and *a* = 0.05 were fixed for all fish and movements.

The modified Izhikevich equations were solved numerically using a fourth-order Runge–Kutta method (RK4).

#### Adding asymmetry

To capture differences between the amplitudes of oscillations on the two sides of the tail, asymmetry was introduced by shifting the xy-oscillations toward either positive or negative values. We used the parameter *shift*, whose magnitude determines the degree of asymmetry and whose sign determines its direction (Figure S3B). *shift* = 0 corresponds to symmetric oscillations, whereas increasing |*shift*| progressively biases the oscillations toward one side. The shift was introduced gradually at the onset of the xy-oscillations, such that it reached the value *shift* over the first half-cycle of the xy-oscillations:

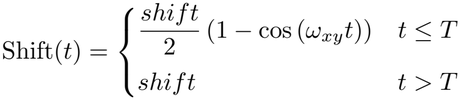

where

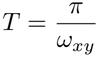

Since *−*1 *≤ shift ≤* 1, Shift(*t*) also remains between *−*1 and 1. Given this time-dependent shift, the xy-dynamics were modified as:

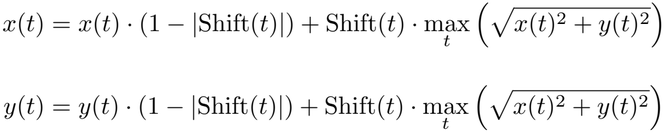

where 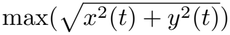 denotes the maximum radial amplitude of the dynamics over time before the shift was introduced. Here, time for the asymmetry transformation was measured relative to the onset of the modeled xy-oscillations.

#### Adjusting oscillation amplitude

To account for variation in the amplitude of the xy-dynamics across movements, the x-dynamics were scaled by *A*_max_. The corresponding y-amplitude was determined using the fish-specific relative amplitude factor *F*_amp_ (Figure S3B,E):

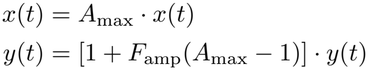

The fish-specific relationship between x- and y-dynamics amplitudes, *F*_amp_, was estimated from movements with an absolute change in heading smaller than 20°, to minimize potential effects of the h-dynamics on this relationship. For each movement, the amplitude of the x- and y-dynamics was defined as half of its peak-to-peak range. We fitted the relationship between x- and y-amplitudes using robust Huber regression with an intercept, and the fitted slope defined the fish-specific relative amplitude factor *F*_*amp*_. Across fish, *F*_*amp*_ was 0.57 *±* 0.09 (mean*±*SD).

Thus, *A*_max_ controlled the overall amplitude of the xy-dynamics, and consequently, the maximal tail curvature, whereas *F*_amp_ preserved the characteristic relationship between x- and y-amplitudes for each fish.

#### Adding decay to return the xy-dynamics to baseline

For movements with substantial asymmetry (|*shift*| *>* 0.05), the oscillations did not naturally return to zero when the oscillatory dynamics terminated. We therefore added an exponential decay to the oscillations, beginning at the last peak of the y-dynamics (*τ*_*dec*_). The decay rate was determined from the modeled y-dynamics according to

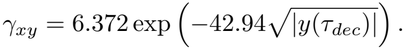

and was applied as follows:

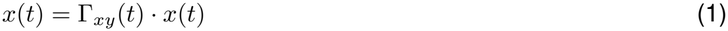

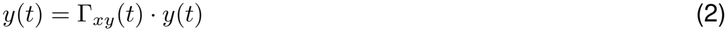

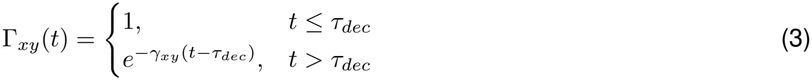

For movements with |*shift*| =*<* 0.05, no additional decay was applied because the dynamics naturally returned close to zero. Because *γ*_*xy*_ was determined entirely from the modeled dynamics, it was not treated as an additional fitted parameter.

Together, the xy-model contained four main parameters describing movement dynamics: *u*_*str*_ (duration), *ω*_*xy*_ (tail-beat frequency), *A*_max_ (maximal tail curvature), and *shift* (asymmetry). *u*_*time*_ and *z*_0_ specified the temporal alignment and phase of the modeled oscillations rather than the movement dynamics themselves.

#### Fitting the xy-dynamics

The parameters of the xy-dynamics model were optimized based on the *R*^2^ between the modeled and observed xy-dynamics, calculated after concatenating the x- and y-dynamics over the fitting window. The fitting window began at the detected movement onset or at the onset of the modeled xy-dynamics, whichever occurred earlier. Modeled onset was defined as the first time point at which the absolute change in either the x- or y-dynamics exceeded 10^*−*9^ before SVD normalization. The end of the fitting window was defined as 20 ms (10 frames) after the last peak of the y-dynamics, corresponding to the onset of the decay defined above, in either the model or the data, whichever occurred later. Thus, modeled oscillations extending beyond the observed movement were penalized by a decrease in fitting *R*^2^.

Fitting was performed hierarchically. We first generated several candidate values of *ω*_*xy*_. Candidate frequencies were obtained from spectral analyses of both the x- and y-dynamics, performed either on the detected movement alone, or on traces containing the detected movement together with the 30 preceding and following frames, with a Tukey window equal to 1 during the detected movement and tapered across the flanking 30-frame intervals. For each analysis, the frequency with maximal power was included as a candidate. An additional candidate frequency was estimated directly from the peak-to-peak interval of the y-dynamics, providing a more accurate estimate at lower frequencies compared to spectral analysis. All candidate frequencies were converted from Hz to angular frequency *ω*_*xy*_ = 2*πf* for use in the model.

For each candidate *ω*_*xy*_, *z*_0_ was estimated from the phase of the corresponding frequency component in the data. We then optimized *u*_*str*_ using SciPy’s *minimize scalar*. For each tested value of *u*_*str*_, *u*_*time*_ was estimated by cross-correlation between the xy-envelope of the data and the model (defined as 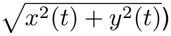, and *shift* was optimized using SciPy’s *minimize scalar*. This procedure provided an estimate of the best fit achievable for each candidate *ω*_*xy*_ and its best *u*_*str*_ value. The frequency yielding the highest fitting *R*^2^ was selected, together with its corresponding *u*_*str*_.

After selecting *ω*_*xy*_ and *u*_*str*_, *u*_*time*_ was first estimated from the xy-envelope of the data and the model using cross-correlation as defined above. It was then refined at a finer temporal resolution based on the *R*^2^ of the full x- and y-dynamics using SciPy’s *minimize scalar* to maximize the fitting *R*^2^. Finally, *shift* was again optimized using SciPy’s *minimize scalar* with all other parameters fixed. During the initial fitting stages, the decay rate *γ*_*xy*_ and its starting point *t*_*decay*_ were fitted directly from the data using Scipy’s *find peaks* and *curve fit* functions, because inaccuracies in the modeled y-dynamics could otherwise bias the model-derived decay rate, which depends on the amplitude at the last y-peak. Once *u*_*str*_ and *u*_*time*_ were fixed, the model-derived *γ*_*xy*_ described above was used. Throughout the fitting, *A*_max_ was calculated directly for each generated model as the non-negative scalar minimizing the squared difference between the modeled and observed x-dynamics:

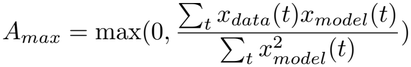

where *x*_*model*_ denotes the modeled x-dynamics before multiplication by *A*_max_. The amplitude of the y-dynamics was then determined from *A*_*max*_ using the fish-specific *F*_*amp*_ described above.

#### Evaluating model-fit accuracy

After fitting the cleaned h-dynamics, the xy-linked component predicted from the modeled x- and y-dynamics was reconstructed using the fish-specific filters described above and added back to obtain the full modeled h-dynamics. Reconstruction accuracy was then quantified as the *R*^2^ between the modeled and original observed dynamics after concatenating the x-, y-, and h-dynamics of each movement. Importantly, when each of the x-, y-, and h-dynamics was concatenated across all movements, their L2 norms differed by less than 2%, indicating that the three dynamics had comparable overall magnitudes and therefore contributed comparably to the reconstruction *R*^2^. This reconstruction *R*^2^ is reported throughout the paper.

### Modeling coupled dynamics

In most movements, the h- and xy-dynamics were modeled independently, such that the time to bend of the h-dynamics and the tail-beat frequency of the xy-dynamics were fitted independently. However, in some movements the two dynamics were coupled: during the bend of the h-dynamics, the tail-beat frequency of the xy-dynamics was determined by the time to bend of the h-dynamics (Figure S4A). For these movements, the xy-dynamics had two values of *ω*_*xy*_: a coupled value, *ω*_*xy*_ = *ω*_*h*_, during the coupled period, and an independently fitted value outside this period. During this period, the absolute amplitude of the y-dynamics was truncated at a fixed value across all coupled movements and fish (0.005 before multiplication by the SVD scaling factor):

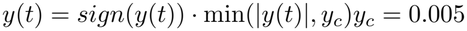

This truncation was observed only in the y-dynamics and was therefore used to define the orientation of the x- and y-axes within the xy-plane, as described above.

The coupled period extended from the onset of the bend to the point in time where |*y*(*t*) | dropped below the trun-cated value (|*y*(*t*) | *< y*_*c*_). At this point, the dynamics were modeled as uncoupled by changing the value of *ω*_*xy*_ in the differential equation to its uncoupled value. To simplify the fitting procedure, we did not model movements in which the y-dynamics are both truncated by the coupled regime and shifted by movement asymmetry. Accordingly, *shift* was fixed at zero for coupled movements. Consequently, neither asymmetry nor the associated end-of-movement decay was included in the coupled model.

#### Fitting coupled dynamics

The coupled dynamics were fitted by first assigning initial values to the three fitted parameters, *ω*_*h*_, *ω*_*xy*_ and *u*_*str*_, and then optimizing them sequentially using SciPy’s *minimize scalar*. For each tested combination of these parameters, the h-dynamics were fitted, with *ω*_*h*_ held fixed and *h*_*delay*_ kept within 5 milliseconds of the xy-dynamics onset. The initial value of *u*_*time*_ was 44.4 ms and was subsequently adjusted based on the cross-correlation between the modeled and observed dynamics.

The initial value of *u*_*str*_ was estimated from the empirical relationship between the integral over the xy-dynamics envelope (Figure 2B) and *u*_*str*_, which was approximately linear for a given *ω*_*h*_. The initial value of *ω*_*h*_ was taken from the independently fitted h-dynamics. Because the xy-frequency changed during the coupled period, *ω*_*xy*_ could not be reliably initialized from spectral analysis. Instead, three candidate frequencies spanning the observed frequency range (24, 31, and 38 Hz) were converted to angular frequencies using *ω*_*xy*_ = 2*πf* and used to initialize three separate coupled-model fits. The frequency with the highest *R*^2^ was retained as the initial guess for the parameter. *z*_0_ was not fitted, but instead fixed at movement onset based on the sign of *h*_*peak*_: *z*_0_ = 5.3 [*rad*] for negative *h*_*peak*_ and *z*_0_ = 7.942 [*rad*] for positive *h*_*peak*_.

Each movement was fitted using both the coupled and uncoupled models, and model assignment was determined solely by which model produced the higher *R*^2^. This automatic model-selection procedure selected coupled fits predominantly for movements with large maximum tail bends (Figure S4B).

### Relating model parameters to movement kinematic features

To describe movements using intuitive kinematic quantities rather than the mathematical parameters of the dynamical model, the fitted parameters were converted to six movement features: maximal tail bend, time to bend, movement duration, tail-beat frequency, maximal tail curvature, and asymmetry.

For the h-dynamics, the signed peak excursion of the modeled dynamics, 2 *\* h*_*peak*_ was used to quantify maximal tail bend. Time to bend was quantified as 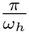, corresponding to the time from bend onset to the peak of the cosine component (Figure S3A).

For the xy-dynamics, increasing the input to the u-dynamics, *u*_*str*_, increased the duration of the transient oscillations. We therefore converted *u*_*str*_ to movement duration by detecting the onset and end of the modeled oscillations and fitting the relationship between the resulting duration and *u*_*str*_:

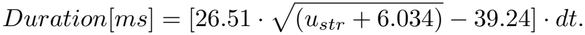

where *dt* = 2 ms is the sampling interval at 500 Hz. This relationship explained 91% of the variance in the modeled movement duration (Figure S3D) and was therefore used to convert *u*_*str*_ into movement duration.

In the modified Izhikevich model, increasing *u*_*str*_ affected both the duration and amplitude of the oscillatory dynamics, such that longer oscillatory episodes also had larger amplitudes before scaling by *A*_max_. Consequently, the *A*_max_ required to produce a given oscillation amplitude depended on *u*_*str*_. We therefore defined maximal tail curvature directly from the amplitude of the modeled x-dynamics rather than from either parameter alone:

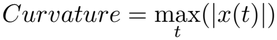

As described above, this quantity was scaled across fish using the second singular value of the SVD decomposition. We verified that maximal tail curvature reflected the joint contributions of *u*_*str*_ and *A*_max_ using linear regression (Figure S3E). Finally, *shift* was used directly to quantify movement asymmetry, while *ω*_*xy*_ was converted to tail-beat frequency by dividing it by 2*π*.

### Model performance across fish

To assess model performance across fish and movements, we first excluded movements for which the extracted tail dynamics were movements with unreliable tail tracking, noisy extracted dynamics, or compound movements that could not be described as a single bout. Tracking reliability was assessed from the extracted tail length, measured between the swim bladder and the tail tip. A movement was considered unreliable if its median tail length deviated by more than 2 MADs (median absolute deviation) from the median tail length of that fish. Movements showing non-smooth changes in the coefficients time series were considered noisy and excluded following manual inspection. Compound movements were likewise identified manually and excluded from the analysis.

In the 5-7 dpf dataset, of the 7,713 detected movements, 681 were excluded based on the tail-length criterion, 277 were excluded as noisy, and 197 were identified as compound movements. Of the remaining 6,558 movements, 6,115 (93%) achieved a reconstruction accuracy of *R*^2^ *>* 0.7. The 13-15 dpf dataset was used to evaluate model generalization and the model-free prediction of movement parameters in a fully automated workflow. Accordingly, no manual screening for noisy or compound movements was performed. Of the 5,006 detected movements, 430 were excluded based on the tail-length criterion, leaving 4,576 movements, of which 4,070 (89%) achieved a reconstruction accuracy of *R*^2^ *>* 0.7.

### Quantifying movement outcomes

Movement outcome was quantified using three parameters: movement distance (*d*), change in heading angle (Δ*θ*_*head*_), and azimuthal angle (*ϕ*_*az*_) (Figure 1B). Larval position was defined as the midpoint between the centers of the two detected eyes (Figure 1B). Movement distance was calculated as the Euclidean distance between the pre- and post-bout larval positions (Figure 1B). The azimuthal angle was defined as the angle between the pre-bout heading direction and the direction of displacement during the bout, with the latter defined by the vector connecting the pre- and post-bout larval positions (Figure 1B). The change in heading angle was defined as the angular difference between the pre- and post-bout heading directions (Figure 1B). For both the change in heading angle and the azimuthal angle, positive values corresponded to a clockwise direction relative to the pre-bout heading, and angle values ranged from *−*180° to 180°.

### Assignment of conventional bout types

To compare the continuous dynamical description with conventional discrete movement categories, bouts were assigned to established zebrafish bout types using the Megabouts classification pipeline ^30^. The data were converted to the input format required by Megabouts, consisting of the spatial x and y coordinates of the head and 11 equidistant points along the tail. Bout types were displayed according to the ordering scheme defined by Megabouts classification.

For prediction of movement parameters from a given bout type (Figure S7L), we performed linear regression using Matlab’s fitlm function, with bout type treated as a categorical variable. This analysis included movements with a model reconstruction *R*^2^ *>* 0.7. For prediction of movement outcomes from bout type (Figure 4G), we restricted the analysis to movements with a model reconstruction *R*^2^ *>* 0.9, consistent with the inclusion criterion used for the corresponding predictions of movement outcome from model parameters (Figure 3F-H). Bout type was again treated as a categorical variable, and linear mixed-effects models were fitted using MATLAB’s fitlme function with fish identity included as a random intercept. The reported values correspond to the adjusted *R*^2^ values returned by the respective models.

### Relating model parameters to movement outcome

To determine how variation in tail dynamics relates to movement outcome (Figure 3F-H), we quantified the relationships between the fitted movement parameters and the resulting changes in fish position and orientation (Δ*θ*_*head*_, *d, ϕ*_*az*_). Unless otherwise stated, these analyses were restricted to movements with a highly accurate model fit *R*^2^ *>* 0.9, ensuring that fitted movement parameters were estimated with high confidence (2588 movements, 39% of the data). Importantly, the relationships between movement parameters and movement outcomes remained strong when the less stringent model-fit criterion of *R*^2^ *>* 0.7 was used: the adjusted *R*^2^ values for change in heading angle, movement distance and azimuthal angle were 0.84, 0.85, and 0.82, respectively, compared with 0.92, 0.91, and 0.90 using the *R*^2^ *>* 0.9 criterion (Figure 3F-H). Relationships between movement parameters and behavioral outcome were quantified using linear mixed-effects models implemented with MATLAB’s fitlme function, with fish identity included as a random intercept to account for inter-fish variability. Predictive performance was evaluated using two-fold cross-validation: the movements of each fish were divided into two groups, a linear mixed-effects model was fitted to one group and used to predict outcomes in the other, and the procedure was then reversed. All reported *R*^2^ values represent the proportion of variance explained in the held-out data of both groups.

To evaluate the contribution of model parameters to movement outcome (Figure 3F-H, S5D-I, S8A), we used reduced models containing the movement parameters that accounted for most of the model performance for each outcome variable. The contribution of additional parameters was assessed by gradually adding them to the model and examining the resulting improvement in adjusted *R*^2^, up to the full parameter model (Figure S5). Change in heading angle was modeled from maximal tail bend, using terms up to third order. Movement distance was modeled from maximal tail bend, including a quadratic term, together with movement duration, maximal tail curvature, and their interaction. Azimuthal angle was modeled from maximal tail bend and movement asymmetry:

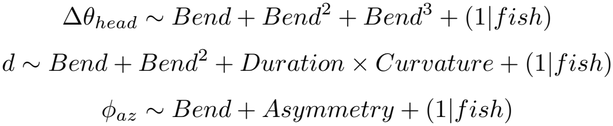

For the azimuth analysis (Figure 3H), maximal tail bend was included as a linear term, as adding higher-order bend terms produced only a modest improvement in adjusted *R*^2^ performance. For the corresponding supplementary analysis (Figure S5I), bend was modeled using terms up to third order:

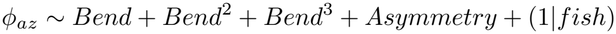

This did not alter the qualitative conclusion that azimuthal angle was primarily captured by maximal tail bend together with movement asymmetry.

To link the six model parameters to movement outcome, we used the following linear mixed models. For the change in heading angle (Figure S5D):

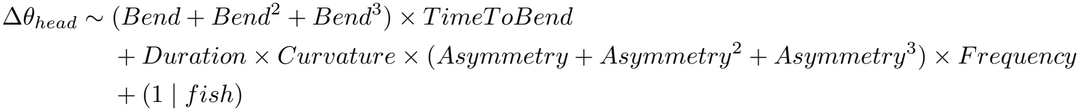

For movement distance (Figure S5E,G):

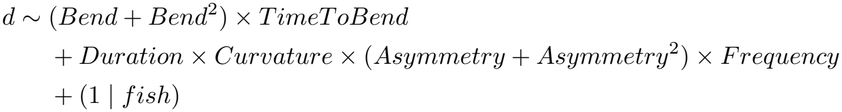

and for azimuthal angle (Figure S5I):

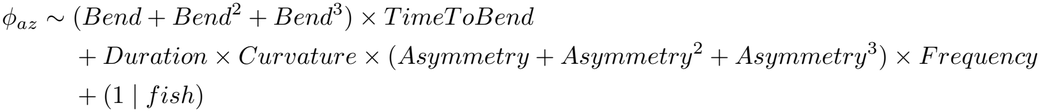

### Relating movement outcome to model parameters

To examine the inverse mapping from movement outcome to tail dynamics (Figure 3I–K), maximal tail bend was predicted from change in heading angle using a third-order polynomial, maximal tail curvature was predicted from log-transformed movement distance, and asymmetry was predicted jointly from change in heading angle and azimuthal angle:

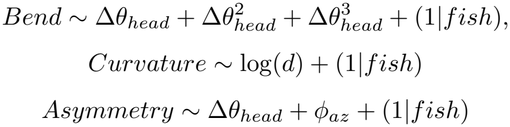

To link asymmetry with movement outcome, coupled movements were not used since we modeled them with *shift* = 0, leaving 2063 movements for analysis. These analyses were restricted to movements with a highly accurate model reconstruction *R*^2^ *>* 0.9. With the less stringent reconstruction criterion of *R*^2^ *>* 0.7, the adjusted *R*^2^ values for maximum tail bend, maximum tail curvature, and asymmetry were 0.86, 0.72, and 0.73, respectively.

To determine how well the three movement outcomes jointly accounted for variation in each model parameter, we additionally fitted models incorporating change in heading angle, azimuthal angle, and movement distance (Figure S6A-B, S8B). For duration, tail-beat frequency, maximal tail curvature, and time to bend, we used a model containing linear and quadratic terms for change in heading angle and azimuthal angle, log-transformed movement distance, and all interactions among these terms

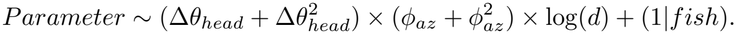

For maximal tail bend and asymmetry, for which the sign carries directional information, we used the signed change in heading angle and azimuthal angle without quadratic terms:

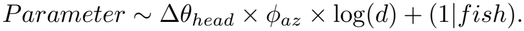

### Model-free prediction of movement parameters from tail movement

To obtain estimates of movement parameters directly from measured tail movements, without fitting the dynamical model to each movement, we developed a model-free prediction procedure. The procedure first projected each tail posture onto a fixed population-average posture basis to obtain the x-, y-, and h-dynamics and then estimated movement parameters directly from features of these dynamics. Thus, once the population-average basis and parameter mappings had been established, movement parameters could be approximated from individual movements without either performing SVD on a large collection of movements or running the full dynamical model-fitting procedure.

To construct the fixed posture basis, we took advantage of the high similarity between the principal posture components across fish (Figure S1B). For each of the 30 fish in the 5–7 dpf dataset, the first three principal posture components were transformed into its cylinder-aligned x, y and h basis (Figure 1G), and the resulting cylinder-aligned posture basis vectors were then averaged across fish. This yielded a population-average posture basis associated with each of the x-, y-, and h-dynamics.

For each tail posture within a movement, we projected each posture onto the three population-average cylinder-aligned basis vectors to obtain the corresponding x, y, and h coefficients. Applying this projection to each frame of a movement yielded the x-, y-, and h-dynamics as time series. We next identified features of these dynamics that could be used to approximate the corresponding model parameters.

Since the maximal tail bend was associated with the most extreme value of the h-dynamics (Figure 2G), the approximated maximal tail bend was defined from the extreme side of the h distribution across the recorded movement: the 90th percentile was used when the positive h extreme was larger than the negative extreme, and the 10th percentile otherwise. Percentiles rather than extrema were used to reduce sensitivity to isolated noisy values:

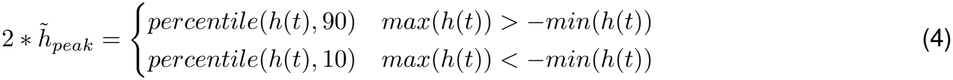

Since maximal tail curvature was mostly reflected in the amplitude of the xy-dynamics (Figure 2I), it was approximated from half the peak-to-peak amplitude of the x-dynamics 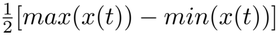.

Because movement asymmetry produced a sustained bias of the x-dynamics toward one side (Figure 2L), it was approximated from the temporal sum of the x-dynamics, normalized by the estimated curvature.

Because these empirical measures provide approximate estimates of the fitted model parameters, their scale and offset could differ systematically from those obtained with the full model-fitting procedure. We therefore used the 5-7 dpf dataset of 30 fish to fit linear relationships between each empirical estimate and its corresponding fitted model parameter. Following this calibration, the final parameter approximations were:

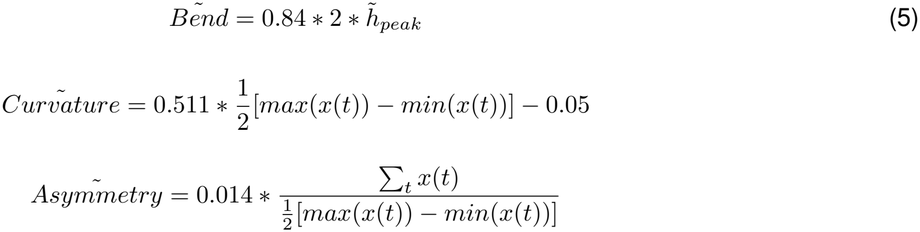

The asymmetry mapping was evaluated for movements with small changes in head orientation (Δ*θ*_*head*_ *<* 20°, 2018 movements, 78% of movements with *R*^2^ *>* 0.9), which were predominantly fitted by the uncoupled model and therefore had a fitted asymmetry parameter.

We next asked whether the same empirical estimates could be used to predict movement outcome. Using the 5-7 dpf dataset, we fitted relationships between these estimates and each movement outcome:

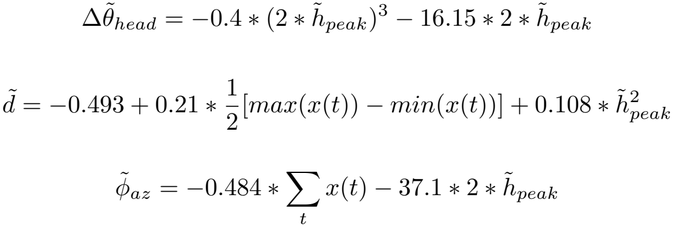

The relationships established in the 5-7 dpf dataset were then tested on the independent 13-15 dpf dataset of 25 fish (Figure 5B–C, Figure S8C–F). These analyses were restricted to movements with a highly accurate model fit *R*^2^ *>* 0.9, ensuring that fitted movement parameters were estimated with high confidence (1700 movements, 37% of the data).

## Acknowledgments

We thank Claire Wyart, Leonard Constien, and Roy Harpaz for their comments on earlier versions of the article. We are also grateful to Yuval Hart, Roi Holzman, Haim Sompolinsky, Israel Nelken, and the Avitan lab members for inspiring and stimulating discussions. This work began as a collaboration with the late Naftali Tishby, whom we remember with admiration. We gratefully acknowledge funding from Israel Science Foundation grant 1684/20.

