## Supplementary Info for "Two co-occurring dynamics explain the organization of a natural movement repertoire"

### Supplementary information

Yoav Rubinstein[1], Maayan Moshkovitz[1], Shai Tishby Tamari[1], Stas Tiomkin[2], Lilach Avitan*[1]
[1]Edmond and Lily Safra Center for Brain Sciences, The Hebrew University of Jerusalem, 9190401, Israel.
[2]Computer Science Department, Whitacre College of Engineering, Texas Tech University, TX, 79409, USA.

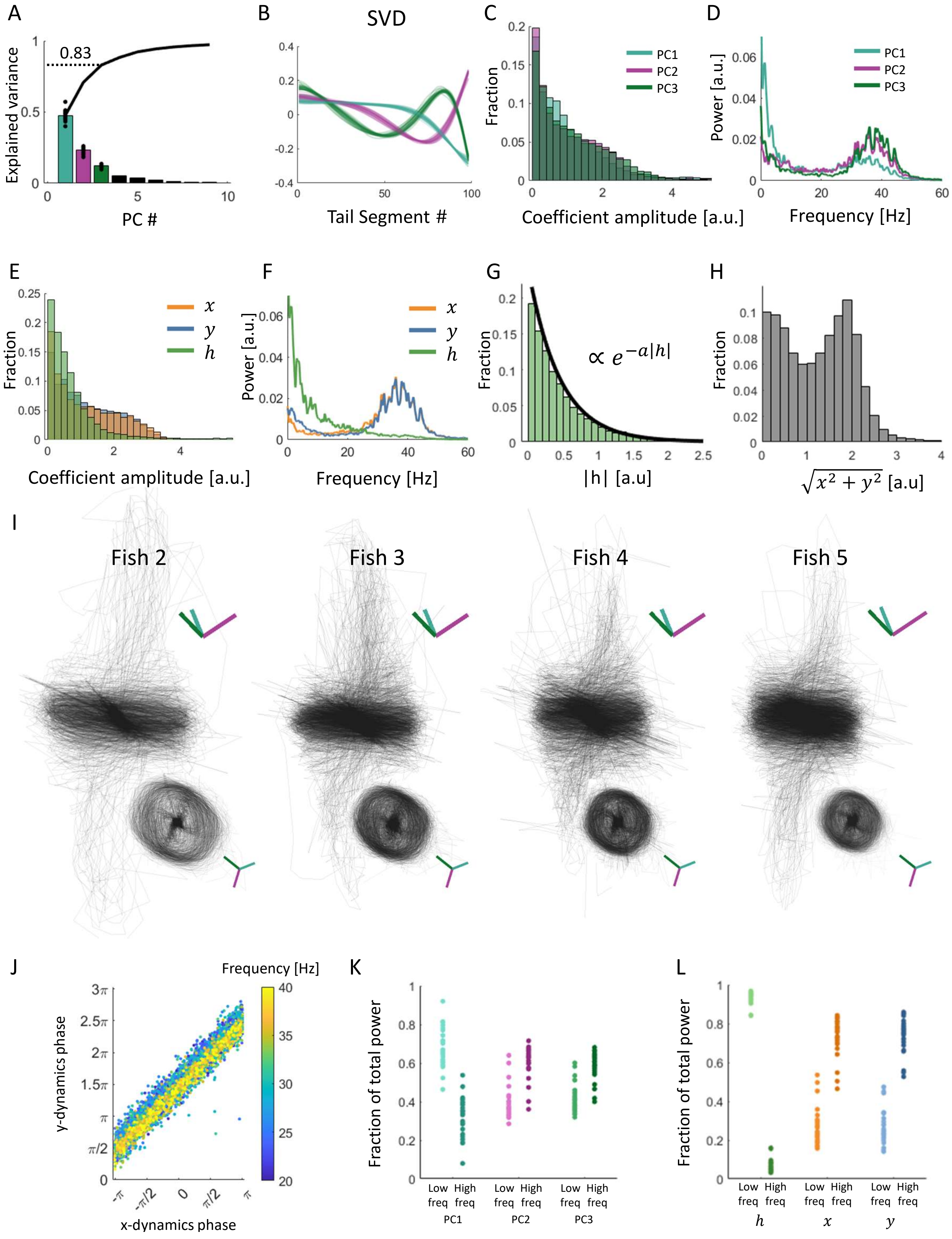
A
Explained variance
0.83
PC #
B
SVD
Tail Segment #
C
Fraction
PC1
PC2
PC3
Coefficient amplitude [a.u.]
D
Power [a.u.]
PC1
PC2
PC3
Frequency [Hz]
E
Fraction
$x$
$y$
$h$
Coefficient amplitude [a.u.]
F
Power [a.u.]
$x$
$y$
$h$
Frequency [Hz]
G
Fraction
$\propto e^{-a|h|}$
|h| [a.u]
H
Fraction
$\sqrt{x^2+y^2}$ [a.u]
I
Fish 2
Fish 3
Fish 4
Fish 5
J
Frequency [Hz]
y-dynamics phase
x-dynamics phase
K
Fraction of total power
Low freq
High freq
PC1
PC2
PC3
L
Fraction of total power
Low freq
High freq
$h$
$x$
$y$

**Figure S1: The low-dimensional organization of tail movements is conserved across fish. A:** Explained variance of tail postures computed separately for each fish. The black line is the mean cumulative explained variance across fish, indicating that, on average, the first three PCs account for 83% of the variance. **B:** The first three posture PCs were computed independently for each fish and showed highly similar shapes across animals, indicating a conserved low-dimensional posture basis. To quantify this similarity, we projected each fish's posture data onto the PCs of a representative fish and compared the resulting PC coefficients with those obtained using the fish's own PCs. The two sets of coefficients showed high agreement ($R^2 = 0.983 \pm 0.015$, mean±SD across fish). **C:** Distribution of the PC1-3 coefficients for the representative fish shown in Figure 1, showing similar distributions across the three components. **D:** Power spectra of the three coefficient time courses for the representative fish. All three components showed increased power around 40Hz, although this feature was less prominent in the first component. **E:** Distribution of the coefficients in the cylinder-aligned coordinates (x, y, h) for the representative fish, showing similar distributions for x and y and a distinct distribution for h, compared with the PC coefficients before rotation in C. **F:** Power spectra of the cylinder-aligned coefficients for the representative fish. The h-dynamics were dominated by lower frequencies, with less power around 40Hz, whereas higher-frequency content was concentrated in the x- and y-dynamics. This frequency separation was weaker in the original PC coordinates, as shown in D. **G-H:** Quantification of the cylindrical geometry in the cylinder-aligned coordinate system (x, y, h) for the representative fish. The distribution of h values followed an approximately exponential distribution (G), indicating that trajectories were concentrated near the middle of the cylinder, with progressively fewer points at larger heights. The distribution of radius in the xy-plane (H) peaked at $2 < \sqrt{x^2 + y^2} < 3$ range, indicating that the trajectory was confined to a characteristic radius. **I-L:** The cylindrical geometry and frequency separation between axes were conserved across fish. **I:** Movement trajectories in the cylinder-aligned coordinate system (x, y, h) for four additional example fish, obtained using the same cylinder-alignment procedure as in Figure 1. Colored axes indicate the original PC1-3 directions for each fish. Across fish, h-dynamics captured displacement along the cylinder height, whereas xy-dynamics captured rotation within a narrow range of radii, demonstrating conserved cylindrical geometry. **J:** The phase difference between the x- and y-dynamics was approximately constant at $\pi$/2, consistent with a single oscillatory dynamic tracing a circular trajectory in the xy-plane. Each dot represents the x- and y-dynamics phase at a single frequency, calculated from a signal formed by concatenating all movements of a representative fish. Shown are the 5% of frequencies with the highest power above 20Hz. **K-L:** Frequency content of the PC coefficient time courses across fish before (K) and after (L) rotation, separated into low-frequency ($< 20$Hz) and high-frequency ($> 20$Hz) components. The y-axis shows the fraction of power contained within each frequency range relative to the power across all frequencies in the dynamics. Before rotation, PC1-3 contained similar mixtures of low- and high-frequency content. After rotation, lower frequencies predominated in h, whereas higher-frequency activity was concentrated in x and y, indicating greater frequency separation in the cylinder-aligned coordinates across fish compared to the original PCs ($p_{val} < 1e{-}15, 1e{-}16, 1e{-}12$ for the x-, y-, and h-dynamics, respectively, paired t-test).

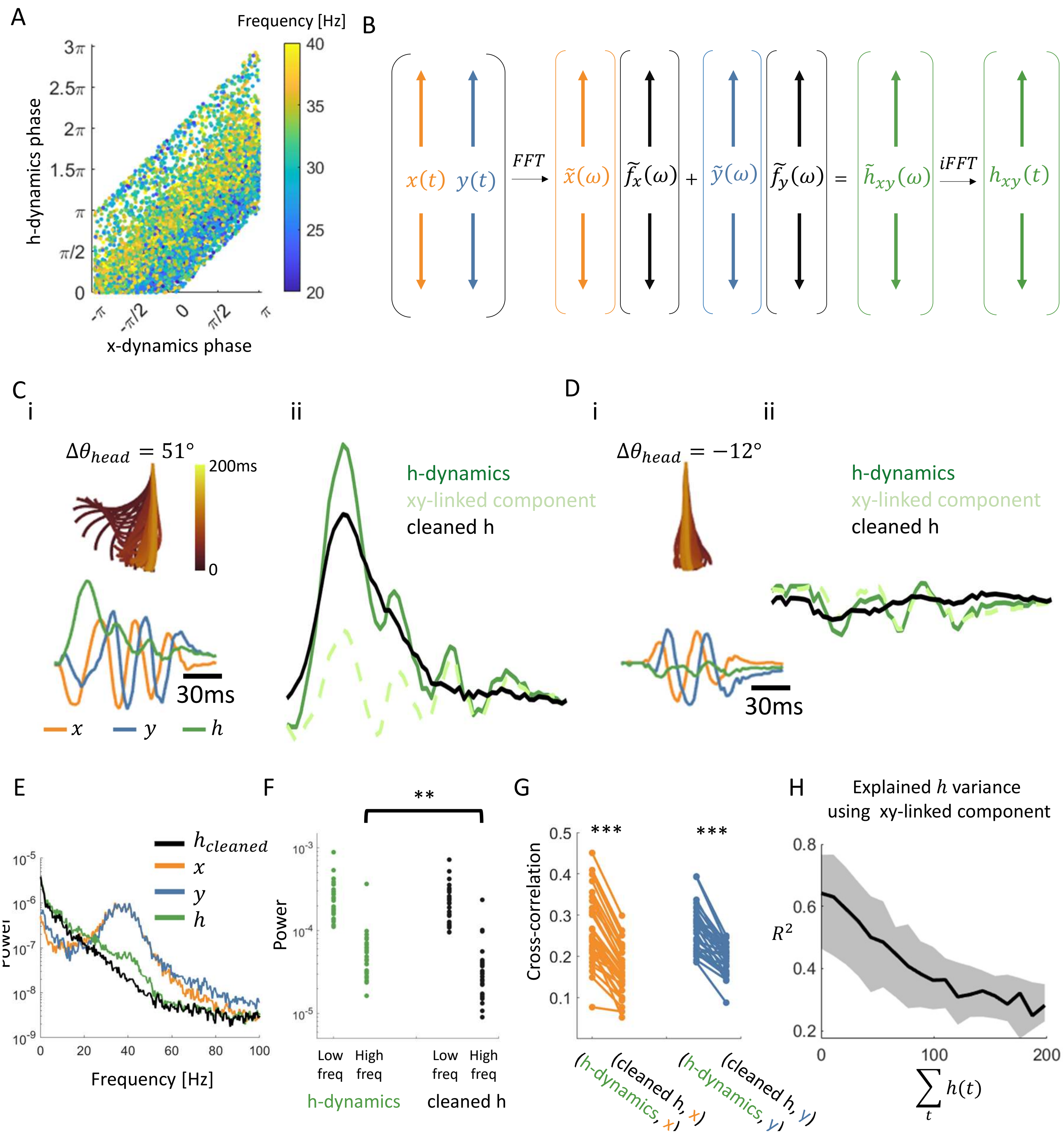
A
Frequency [Hz]
h-dynamics phase
x-dynamics phase
B
FFT
iFFT
C
i
ii
$\Delta\theta_{head} = 51°$
200ms
30ms
h-dynamics
xy-linked component
cleaned h
D
i
ii
$\Delta\theta_{head} = -12°$
30ms
h-dynamics
xy-linked component
cleaned h
E
Frequency [Hz]
F
Power
Low freq
High freq
Low freq
High freq
h-dynamics
cleaned h
G
Cross-correlation
(h-dynamics, x)
(cleaned h, x)
(h-dynamics, y)
(cleaned h, y)
H
Explained $h$ variance using xy-linked component
$R^2$
$\sum_t h(t)$

**Figure S2: Removing a small xy-linked residual from the h-dynamics sharpens the separation between h- and xy-dynamics. A:** After rotation into cylinder-aligned axes, a small high-frequency residual remained in the h-dynamics. Its phase was systematically related to the phases of the x- and y-dynamics, suggesting that it was linked to the xy-dynamics and could therefore be predicted from them. Each dot represents the x- and y-dynamics phase at a single frequency, calculated from a signal formed by concatenating all movements of a representative fish. Shown are the 5% of frequencies with the highest power above 20Hz. **B:** Schematic of the method used to estimate the xy-linked component from x- and y-dynamics. The x- and y- dynamics, $x(t)$ and $y(t)$, were transformed into Fourier space $\tilde{x}(\omega)$ and $\tilde{y}(\omega)$. Fish-specific filters, $\tilde{f}_x(\omega)$ and $\tilde{f}_y(\omega)$, were applied to $\tilde{x}(\omega)$ and $\tilde{y}(\omega)$, respectively, and the filtered signals were summed to reconstruct the xy-linked component of the h-dynamics, ($\tilde{h}_{xy}(\omega)$). This component was then transformed back into the time domain, $h_{xy}(t)$, and subsequently subtracted from the original h-dynamics. The filters $\tilde{f}_x(\omega)$ and $\tilde{f}_y(\omega)$ were derived separately for each fish (see Methods). **C:** Removal of the xy-linked component from the h-dynamics, illustrated for an example movement with a large integral of the h-dynamics over time. (i) Top: tail posture dynamics over time for the example movement; bottom: the corresponding h- and xy-dynamics. (ii) Removing the xy-linked component from the h-dynamics revealed a cleaned h-dynamics characterized by a single rise to a peak followed by a return to baseline. **D:** Removal of the xy-linked component from the h-dynamics, illustrated for an example movement with a small integral of the h-dynamics over time. (i) Top: tail posture dynamics over time for the example movement; bottom: the corresponding h- and xy-dynamics. (ii) Removing the xy-linked component from the h-dynamics revealed a cleaned h-dynamic of small amplitude, again characterized by a single excursion to a peak or trough followed by a return to baseline. This procedure further reduced the correlations between the h-dynamics and the x- and y-dynamics across movements, yielding peak cross-correlations across lags of $r_{x,h} = 0.19$, $r_{y,h} = 0.19$, respectively, compared with the corresponding values before removal of the xy-linked component (Figure 1H). **E:** While the h-dynamics showed a small spectral peak around 40Hz, corresponding to the prominent peaks in the x- and y-dynamics, this peak was removed after subtracting the xy-linked component, as shown by the spectrum of $h_{cleaned}$. Note the logarithmic y-axis, which makes this small change visible (compare with Figure S1F). **F:** Across fish, removal of the xy-linked component reduced the high-frequency ($> 20$Hz) content of the h-dynamics ($p < 1e^{-4}$, paired t-test). **G:** Across fish, removing the xy-linked component reduced the correlation between the h-dynamics and the x- and y-dynamics ($p < 1e^{-11}$ for both dynamics, paired t-test). **H:** The xy-linked component accounted for part of the variance in the h-dynamics. Its relative contribution was greatest in movements with weak h-dynamics, as quantified by the integral of h over the movement, and progressively decreased as this integral increased, as illustrated in C and D. The black solid line shows the median, and the gray shading indicates the 25th-75th percentile of movements in the dataset.

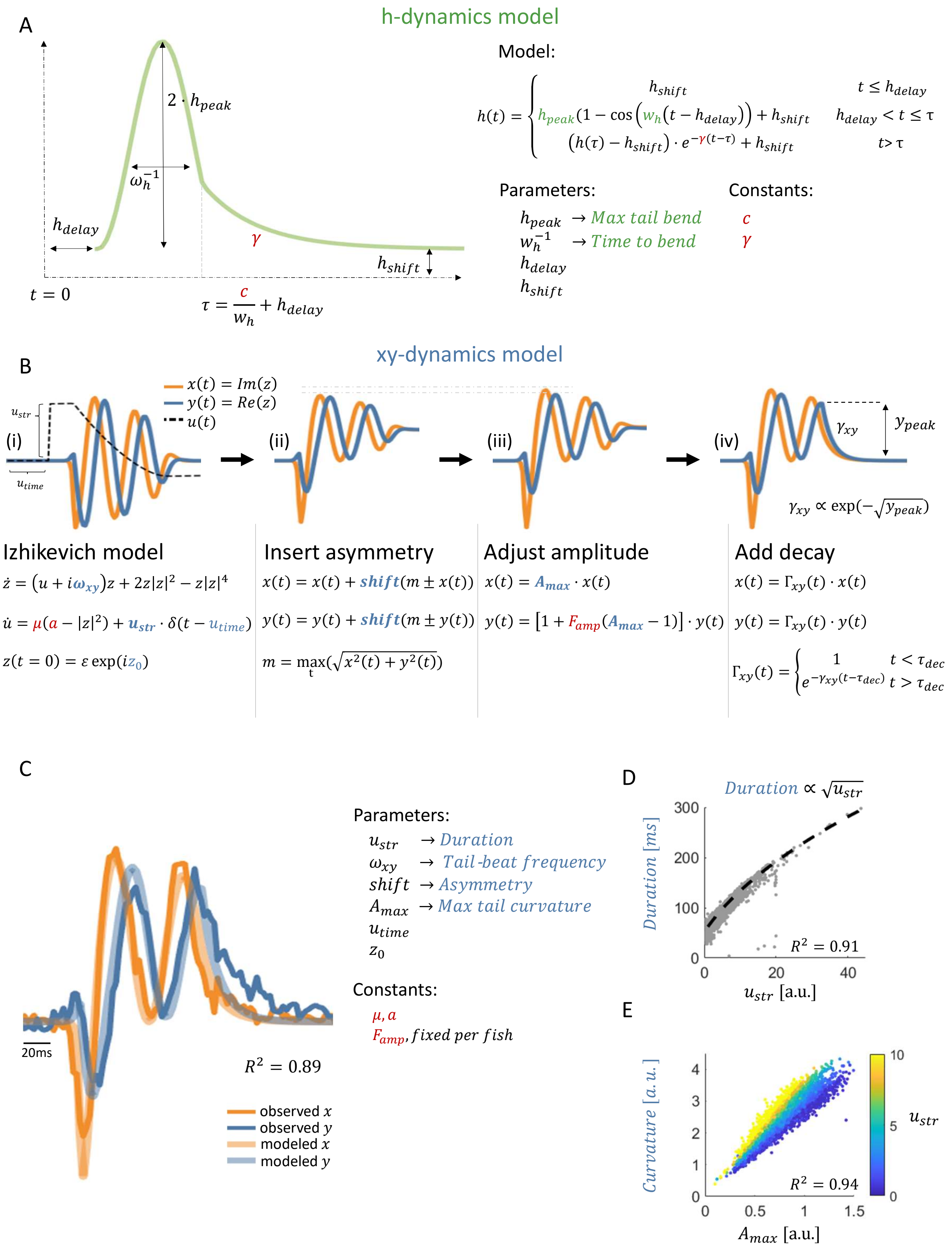
A
h-dynamics model
Model:
$h(t) = \begin{cases} h_{shift} & t \le h_{delay} \\ h_{peak}\left(1-\cos\left(w_h(t-h_{delay})\right)\right) + h_{shift} & h_{delay} < t \le \tau \\ \left(h(\tau)-h_{shift}\right)\cdot e^{-\gamma(t-\tau)} + h_{shift} & t > \tau \end{cases}$
$2 \cdot h_{peak}$
$\omega_h^{-1}$
$h_{delay}$
$\gamma$
$h_{shift}$
$t = 0$
$\tau = \frac{c}{w_h} + h_{delay}$
Parameters:
$h_{peak}$ → Max tail bend
$w_h^{-1}$ → Time to bend
$h_{delay}$
$h_{shift}$
Constants:
$c$
$\gamma$
B
xy-dynamics model
$x(t) = Im(z)$
$y(t) = Re(z)$
$u(t)$
$u_{str}$
$u_{time}$
(i)
(ii)
(iii)
(iv)
$\gamma_{xy}$
$y_{peak}$
$\gamma_{xy} \propto \exp(-\sqrt{y_{peak}})$
Izhikevich model
$\dot{z} = (u + i\omega_{xy})z + 2z|z|^2 - z|z|^4$
$\dot{u} = \mu(a - |z|^2) + u_{str} \cdot \delta(t - u_{time})$
$z(t=0) = \varepsilon \exp(iz_0)$
Insert asymmetry
$x(t) = x(t) + shift(m \pm x(t))$
$y(t) = y(t) + shift(m \pm y(t))$
$m = \max_t(\sqrt{x^2(t) + y^2(t)})$
Adjust amplitude
$x(t) = A_{max} \cdot x(t)$
$y(t) = [1 + F_{amp}(A_{max} - 1)] \cdot y(t)$
Add decay
$x(t) = \Gamma_{xy}(t) \cdot x(t)$
$y(t) = \Gamma_{xy}(t) \cdot y(t)$
$\Gamma_{xy}(t) = \begin{cases} 1 & t < \tau_{dec} \\ e^{-\gamma_{xy}(t-\tau_{dec})} & t > \tau_{dec} \end{cases}$
C
20ms
$R^2 = 0.89$
observed $x$
observed $y$
modeled $x$
modeled $y$
Parameters:
$u_{str}$ → Duration
$\omega_{xy}$ → Tail-beat frequency
$shift$ → Asymmetry
$A_{max}$ → Max tail curvature
$u_{time}$
$z_0$
Constants:
$\mu, a$
$F_{amp}$, fixed per fish
D
$Duration \propto \sqrt{u_{str}}$
Duration [ms]
300
200
100
0
0
20
40
$u_{str}$ [a.u.]
$R^2 = 0.91$
E
Curvature [a.u.]
4
3
2
1
0
0
0.5
1
1.5
$A_{max}$ [a.u.]
$R^2 = 0.94$
10
5
0
$u_{str}$

**Figure S3: Full parameterization of the h- and xy-dynamics model. A:** Parameterization of the h-dynamics after removal of the xy-linked component. The cleaned h-dynamics were modeled as a cosine-shaped rise to a peak, followed by an exponential return to baseline beginning at a constant cosine phase $\tau$. The model included four fitted parameters: max tail bend and time to bend, together with the start time of the modeled dynamics relative to the detected movement onset ($h_{delay}$), and a constant offset $h_{shift}$. The parameters $c$ and $\gamma$ were fixed for all fish and movements. **B:** Full model of the xy-dynamics, constructed in four successive stages (see Methods). (i) Symmetric oscillatory dynamics were modeled using the Izhikevich-inspired oscillator comprising a fast complex variable, $z$, and a latent slow variable $u$. A transient change in $u$ initiated oscillations in $z$ at a frequency $\omega_{xy}$, whereas the amplitude of the transient change in $u$, $u_{str}$, determined their duration (see D). The x- and y-dynamics corresponded to the imaginary and real parts of $z$, respectively. The basic oscillatory model included four fitted parameters: duration and frequency, together with the start time of the modeled dynamics relative to the detected movement onset ($u_{time}$), and the phase of $z$ at that time ($z_0$). Two additional model parameters marked in red, $a$ and $\mu$, affected the dynamics of the latent variable $u$ and were fixed across fish (see Methods). (ii) Asymmetry was then introduced by shifting the oscillation baseline using a parameter ($shift$) constrained to $-1 < shift < 1$. The example shown has pronounced asymmetry, although asymmetry was small in many movements and produced only minor changes. For example, asymmetry is zero in Figure 2B, whereas Figure 2L illustrates the effects of progressively increasing asymmetry. The equations below describe the xy-dynamics following the gradual introduction of asymmetry, as described in details in the Methods. (iii) To account for variations in maximum tail curvature, the model allowed the overall oscillation amplitude to vary while preserving a characteristic relationship between the x- and y- amplitudes. The dashed gray lines indicate the x-and y- amplitudes after amplitude adjustments, allowing comparison with the oscillation amplitudes before the adjustment. This relationship was estimated empirically for each fish, such that setting the x-amplitude also determined the y-amplitude. The relationship was similar across fish ($F_{amp} = 0.57 \pm 0.09$, mean $\pm$ SD). (iv) An exponential decay was then applied to return the oscillatory dynamics to baseline. The decay began at the last significant peak of the y-dynamics identified after stage (iii) and had a decay rate $\gamma_{xy}$ that was determined by the height of that peak. For movements with low asymmetry ($< 0.05$), the dynamics naturally returned close to baseline, and no decay was applied. **C:** Observed (solid lines) and modeled (transparent lines) xy-dynamics for the example movement shown in B, showing a reconstruction of a movement from our dataset with $R^2 = 0.89$. See Methods for the detailed relationship between model parameters and movement features. **D:** Relationship between movement duration and the transient amplitude of the slow variable $u$, $u_{str}$. Increased $u_{str}$ resulted in an increased movement duration such that $u_{str}$ was proportional to its square root. **E:** Once $u_{str}$ was set to produce the specified movement duration, the parameter $A_{max}$ scaled the oscillation amplitude independently of duration, allowing the model to fit maximum tail curvature across movements of different durations.

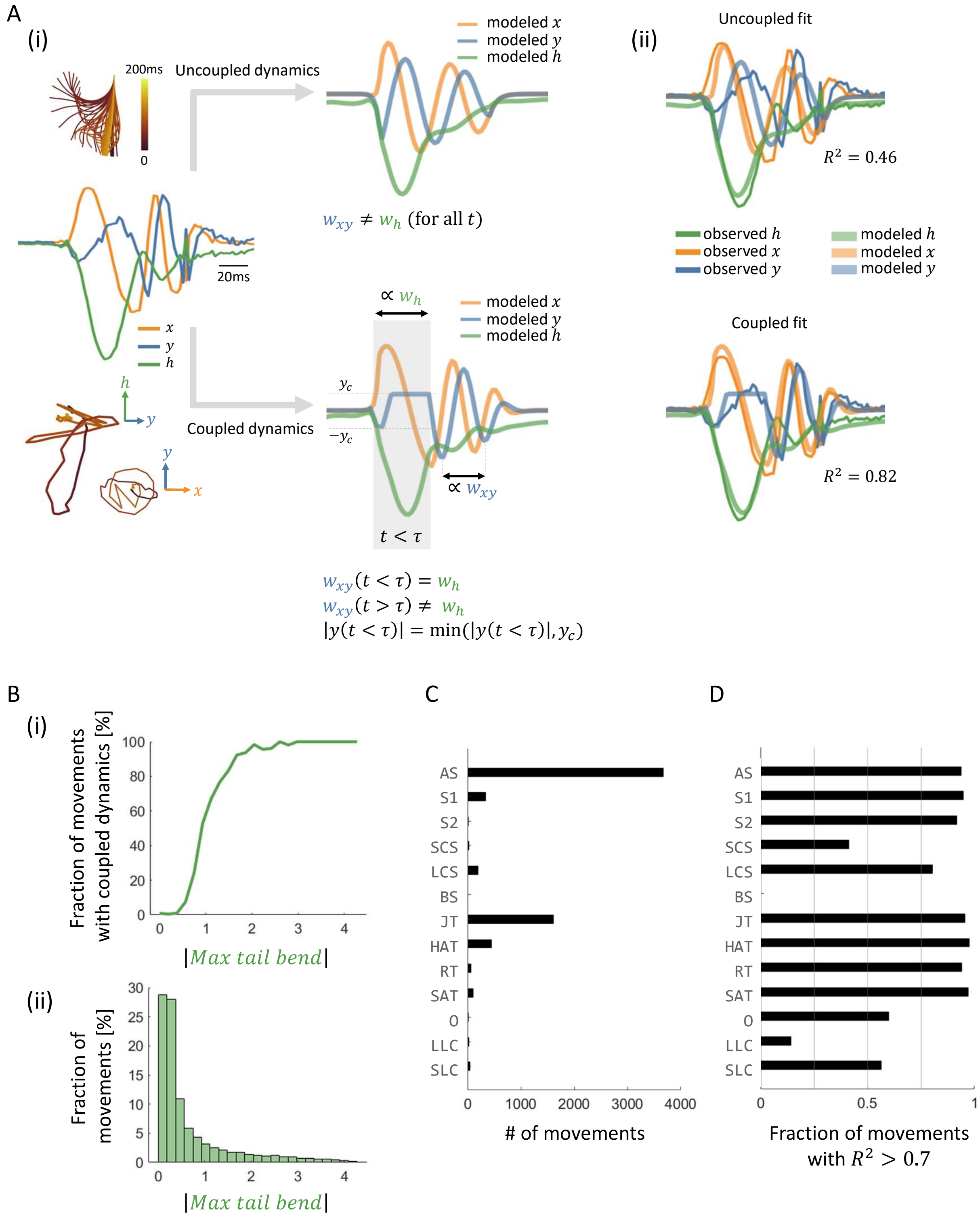

A
(i)
Uncoupled dynamics
200ms
0
modeled x
modeled y
modeled h
20ms
x
y
h
$w_{xy} \neq w_h$ (for all $t$)
$\propto w_h$
Coupled dynamics
$y_c$
$-y_c$
$\propto w_{xy}$
$t < \tau$
$w_{xy}(t < \tau) = w_h$
$w_{xy}(t > \tau) \neq w_h$
$|y(t < \tau)| = \min(|y(t < \tau)|, y_c)$
(ii)
Uncoupled fit
$R^2 = 0.46$
observed h
observed x
observed y
modeled h
modeled x
modeled y
Coupled fit
$R^2 = 0.82$
B
(i)
Fraction of movements with coupled dynamics [%]
|Max tail bend|
(ii)
Fraction of movements [%]
|Max tail bend|
C
AS
S1
S2
SCS
LCS
BS
JT
HAT
RT
SAT
O
LLC
SLC
### of movements
D
Fraction of movements with $R^2 > 0.7$

**Figure S4: Movements with large maximum tail bend showed coupling between the xy- and h-dynamics. A:** Modeling of a movement with large maximum tail bend using coupled dynamics. (i) Top: in the conventional, uncoupled model, the tail-beat frequency of the xy-dynamics remained constant throughout the movement and was independent of the time to bend parameter of the h-dynamics, resulting in a poor fit. Bottom: We therefore introduced coupled dynamics, in which the tail-beat frequency of the xy-dynamics was proportional to the inverse time to bend during a defined epoch encompassing the bend (transparent gray area). During this epoch, the maximum absolute amplitude of the y-dynamics was capped at a fixed value $y_c$, shared across fish and movements. Once the h-dynamics returned to small values, the xy-dynamics resumed an independent tail-beat frequency, and the y-amplitude was no longer capped. Crucially, the duration of the coupled regime was determined by the phase of the h-dynamics, and therefore introduced no additional fitted parameters. (ii) Model fit for the example movement using both methods, coupled (top) and uncoupled (bottom) dynamics, producing a more accurate fit using coupled dynamics indicated by $R^2$ of each fit. **B:** (i) When allowed to fit either coupled or non-coupled dynamics, the model selected the fit with the higher reconstruction $R^2$. Uncoupled dynamics were selected for movements with low maximum tail bend and coupled dynamics for movements with large maximum tail bend. (ii) The distribution of maximum tail bend values indicates that most movements in the dataset had low maximum tail bend. **C-D:** Model performance on 13 bout types identified by Megabouts (See methods). **C:** Number of movements assigned to each bout type in the dataset. **D:** Fraction of movements within each bout type that were reconstructed by the model with $R^2 > 0.7$. Bout types shown are approach swim (AS), slow1 (S1), slow2 (S2), burst swim (BS), short capture swim (SCS), long capture swim (LCS), J-turn (JT), high angle turn (HAT), routine turn (RT), shadow avoidance turn (SAT), O-bend (O), long-latency C-start (LLC) and short-latency C-start (SLC).

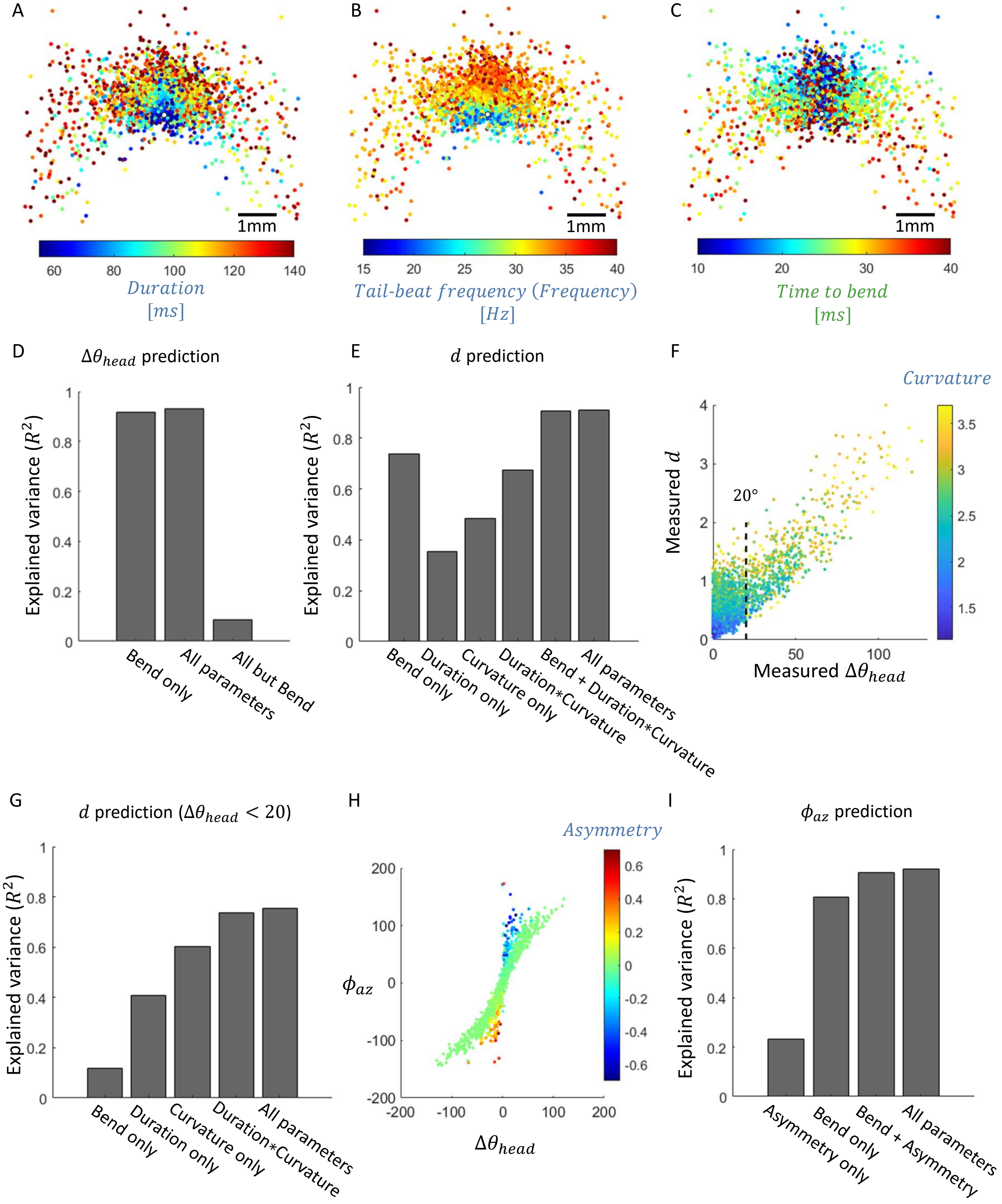
A
1mm
60 80 100 120 140
Duration [ms]
B
1mm
15 20 25 30 35 40
Tail-beat frequency (Frequency) [Hz]
C
1mm
10 20 30 40
Time to bend [ms]
D
Δθ_head prediction
Explained variance (R²)
Bend only
All parameters
All but Bend
E
d prediction
Explained variance (R²)
Bend only
Duration only
Curvature only
Duration*Curvature
Bend + Duration*Curvature
All parameters
F
Curvature
Measured d
Measured Δθ_head
20°
G
d prediction (Δθ_head < 20)
Explained variance (R²)
Bend only
Duration only
Curvature only
Duration*Curvature
All parameters
H
Asymmetry
φ_az
Δθ_head
I
φ_az prediction
Explained variance (R²)
Asymmetry only
Bend only
Bend + Asymmetry
All parameters

**Figure S5: Distinct movement parameters contribute to different aspects of movement outcome. A-C:** Movement outcome repertoire colored by movement duration (A), tail-beat frequency (B), and time to bend (C). Movement duration and tail-beat frequency varied gradually with distance traveled, whereas time to bend showed only a limited relationship to change in heading angle, primarily among large-turn movements. **D:** Cross-validated prediction of change in heading angle, $\Delta\theta_{head}$, from movement parameters. Maximum tail bend alone predicted changes in heading angle with high accuracy ($R^2 = 92$), and including the five remaining movement parameters produced only a modest improvement ($R^2 = 0.93$). By contrast, a model including all parameters except maximum tail bend showed poor predictive performance ($R^2 = 0.09$). **E:** Cross-validated prediction of distance traveled, $d$, from different combinations of movement parameters. Among single-parameter models, maximum tail bend provided the highest predictive performance ($R^2 = 0.74$), followed by maximum tail curvature ($R^2 = 0.48$), and movement duration ($R^2 = 0.35$). Combining maximum tail curvature, duration, and their interaction improved predictive performance ($R^2 = 0.67$). Adding maximum tail bend further increased predictive performance ($R^2 = 0.91$), whereas including the remaining parameters produced no additional improvement ($R^2 = 0.91$). **F:** The relationship between the change in heading angle and distance traveled explained the large contribution of maximum tail bend to variation in distance. Movements are colored by maximum tail curvature. For small changes in heading angle, $(|\Delta\theta_{head}| < 20°)$, distance varied with maximum tail curvature, revealing a curvature gradient across displacements. **G:** Cross-validated prediction of distance traveled for movements with small changes in heading angle. Within this subset, maximum tail bend had low predictive performance ($R^2 = 0.12$), whereas maximum tail curvature was the strongest individual predictor of distance ($R^2 = 0.60$), followed by movement duration ($R^2 = 0.41$). Combining curvature and duration, including their interaction, improved predictive performance ($R^2 = 0.74$), while adding the remaining movement parameters produced only a marginal improvement ($R^2 = 0.75$). **H:** Effect of asymmetry on the relationship between azimuthal angle and change in heading angle. For movements with near-zero asymmetry, azimuthal angle varied approximately linearly with change in heading angle. At larger asymmetry values, this relationship was disrupted, and azimuthal angle further increased. **I:** Cross-validated prediction of azimuthal angle from movement parameters. Maximum tail bend alone provided high predictive performance ($R^2 = 0.81$), reflecting the predominance of approximately symmetric movements in which azimuthal angle closely follows the change in heading angle. Movement asymmetry accounted for deviations from this relationship, improving predictive performance when combined with maximum tail bend ($R^2 = 0.91$). Including the remaining movement parameters produced only a marginal further improvement ($R^2 = 0.92$). All regressions were performed using linear mixed-effects models with fish identity as a random intercept and evaluated using two-fold cross-validation. Maximum tail bend was either treated as a 3rd power polynomial (for $\Delta\theta_{head}$ and $\phi_{az}$ calculations) or as a 2nd power polynomial (for $d$ estimations, see Methods).

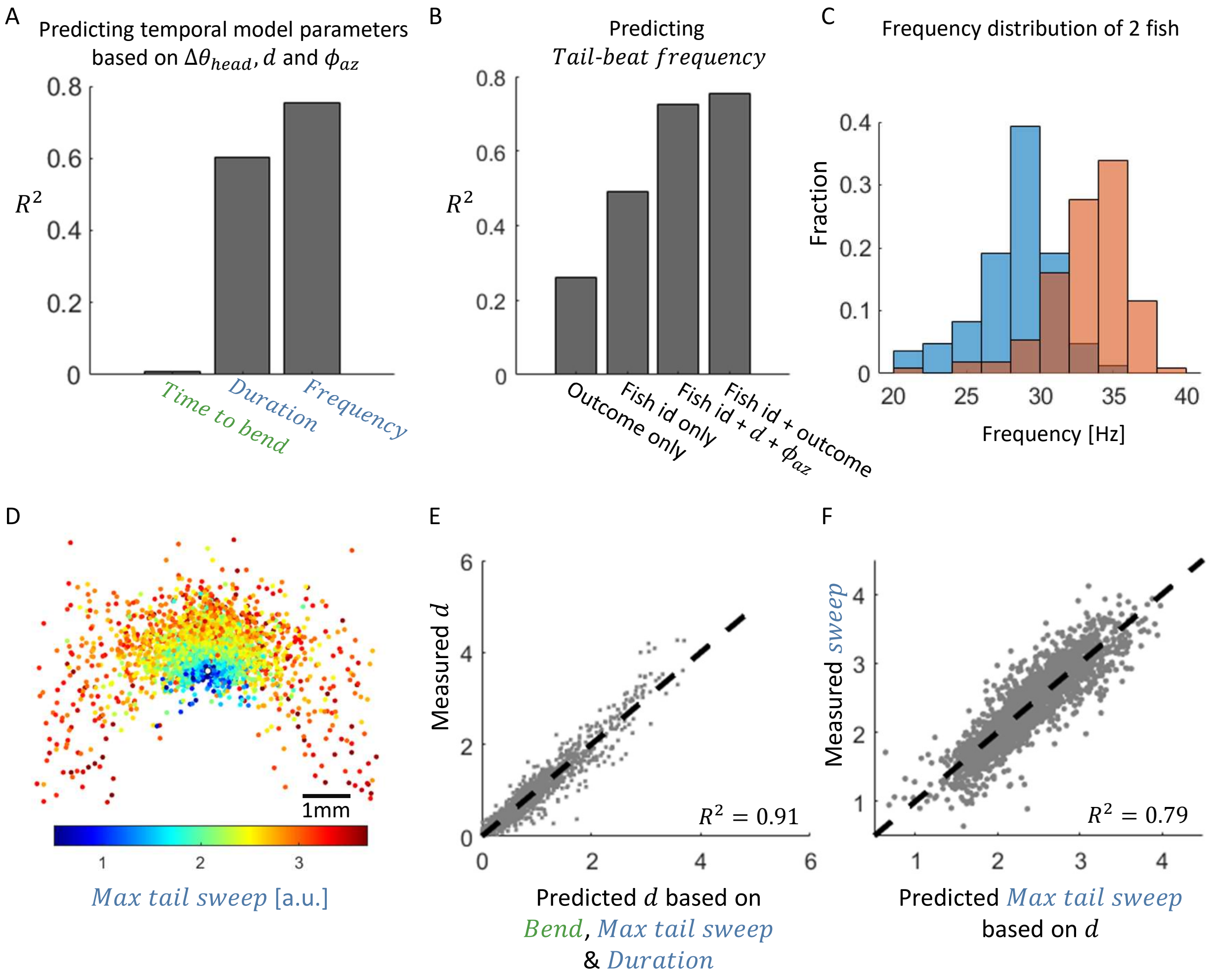

A
Predicting temporal model parameters based on $\Delta\theta_{head}$, $d$ and $\phi_{az}$
$R^2$
0.8
0.6
0.4
0.2
0
Time to bend
Duration
Frequency
B
Predicting Tail-beat frequency
$R^2$
0.8
0.6
0.4
0.2
0
Outcome only
Fish id only
Fish id + $d$ + $\phi_{az}$
Fish id + outcome
C
Frequency distribution of 2 fish
Fraction
0.4
0.3
0.2
0.1
0
20
25
30
35
40
Frequency [Hz]
D
1mm
1
2
3
Max tail sweep [a.u.]
E
Measured $d$
Predicted $d$ based on Bend, Max tail sweep & Duration
$R^2 = 0.91$
F
Measured sweep
Predicted Max tail sweep based on $d$
$R^2 = 0.79$

**Figure S6: Movement temporal parameters are linked to both fish identity and movement outcome. A:** Variation in the movement's temporal parameters (time to bend, movement duration, and tail-beat frequency) was accounted for by the outcome variables ($d$, $\Delta\theta_{head}$, and $\phi_{az}$) ($R^2 = 0.01, 0.6, 0.75$, respectively). Given the outcome variable, explained variance in movement duration and tail-beat frequency was larger than time to bend, consistent with their relationships with movement outcome (see Figure S5A-C). **B:** A modest fraction of the variance in tail-beat frequency was accounted for by the outcome variables alone ($d$, $\Delta\theta_{head}$, and $\phi_{az}$, $R^2 = 0.26$), larger fraction by fish identity ($R^2 = 0.49$), and the largest fraction by their combination ($R^2 = 0.75$). Thus, tail-beat frequency was primarily fish-dependent, but outcome variables explained additional variance beyond fish identity. **C:** Example tail-beat frequency distributions for two fish ($33.7 \pm 2.8$ Hz, $28.3 \pm 2.8$ Hz, mean$\pm$SD). **D-F:** As an additional empirical proxy for maximum tail curvature, we computed the peak-to-peak amplitude of the x-dynamics, defined as the difference between the maximum and minimum x values during the movement and representing the maximum tail sweep. Maximum tail sweep was associated with distance traveled (D). Maximum tail bend, maximum tail sweep, and movement duration together accounted for most of the variance in distance (E), whereas distance accounted for a large fraction of the variance in maximum tail sweep (F).

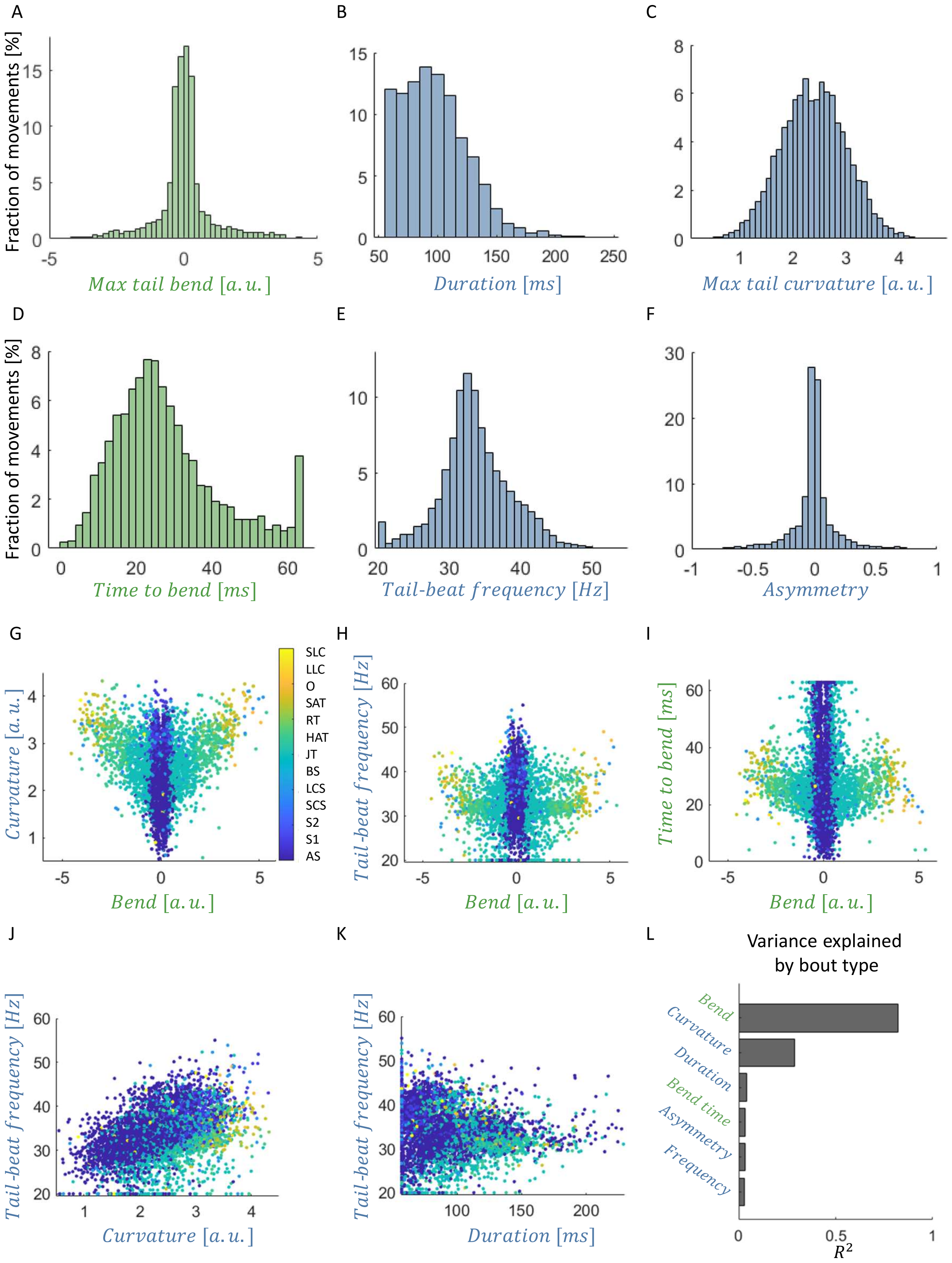

A
Fraction of movements [%]
Max tail bend [a.u.]
B
Duration [ms]
C
Max tail curvature [a.u.]
D
Fraction of movements [%]
Time to bend [ms]
E
Tail-beat frequency [Hz]
F
Asymmetry
G
SLC
LLC
O
SAT
RT
HAT
JT
BS
LCS
SCS
S2
S1
AS
Curvature [a.u.]
Bend [a.u.]
H
Tail-beat frequency [Hz]
Bend [a.u.]
I
Time to bend [ms]
Bend [a.u.]
J
Tail-beat frequency [Hz]
Curvature [a.u.]
K
Tail-beat frequency [Hz]
Duration [ms]
L
Variance explained by bout type
Bend
Curvature
Duration
Bend time
Asymmetry
Frequency
$R^2$

**Figure S7: Movement parameters occupy a structured and constrained parameter space. A-F:** Distributions of the six fitted model parameters across movements reconstructed with a model fit $R^2 > 0.7$. **G:** Relationship between maximum tail bend and maximum tail curvature, showing that large maximum tail bend values were associated with a restricted range of maximum tail curvature values. Movements are colored and ordered by their bout-type assignment. **H:** Relationship between maximum tail bend and tail-beat frequency, showing that a range of tail-beat frequencies was observed at each maximum tail bend value. Because individual fish occupied distinct tail-beat-frequency ranges (Figure S6B,C), the horizontal bands largely correspond to different fish. **I:** Relationship between maximum tail bend and the time to bend, showing that time to bend increased with maximum tail bend for large bend values. **J:** Relationship between maximum tail curvature and tail-beat frequency, which were only weakly related. **K:** Relationship between movement duration and tail-beat frequency, showing a broad range of tail-beat frequencies at a given movement duration. Given the other three xy-dynamics parameters and the interactions between them, tail-beat frequency was predicted with $R^2 = 0.64$, showing that frequency partially correlates with duration, curvature, and asymmetry. **L:** Fraction of variance in each model parameter explained by bout type. Bout type explained the largest fraction of variance in maximum tail bend, followed by maximum tail curvature and movement duration. Thus, bout-type organization aligned most strongly with variation along the bend axis, while also reflecting variation in other movement parameters.

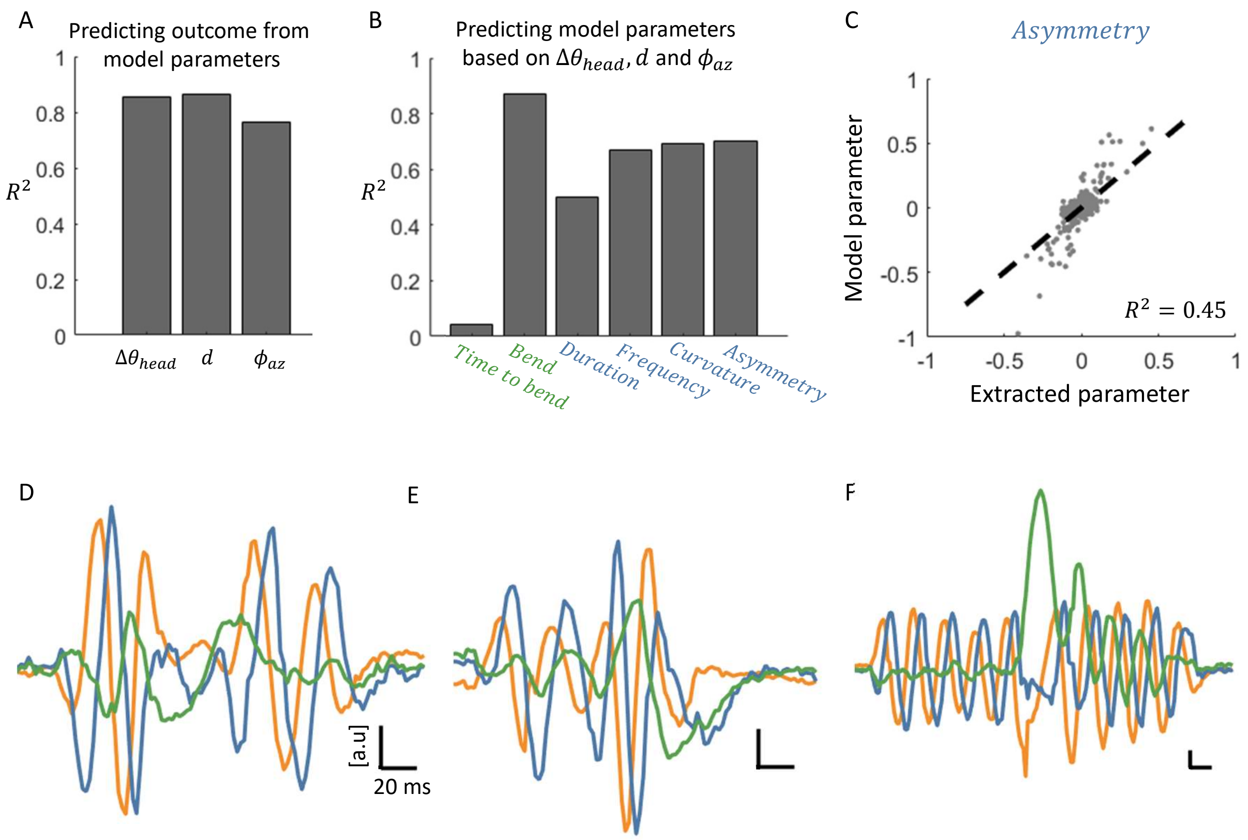
A
Predicting outcome from model parameters
$R^2$
1
0.8
0.6
0.4
0.2
0
$\Delta\theta_{head}$
$d$
$\phi_{az}$
B
Predicting model parameters based on $\Delta\theta_{head}$, $d$ and $\phi_{az}$
$R^2$
Time to bend
Bend
Duration
Frequency
Curvature
Asymmetry
C
Asymmetry
Model parameter
Extracted parameter
$R^2 = 0.45$
D
E
F
[a.u]
20 ms

**Figure S8: The dynamical framework generalizes across development and reveals compound movements. A:** Explained variance of individual movement outcome variables in the 13-15 dpf fish dataset based on the model parameters. All three outcome parameters were accurately predicted, with performance comparable to that obtained in the 5-7 dpf dataset ($R^2 = 0.85, 0.86$ and $0.77$ for $\Delta\theta_{head}, d$ and $\phi_{az}$, respectively. Compare to Figure 3F-H). **B:** Explained variance of individual movement parameters in the 13-15 dpf fish dataset based on outcome variables. Performance was comparable to that obtained in the 7 dpf dataset ($R^2 = 0.04, 0.87, 0.5, 0.66, 0.69$ and $0.69$ for time to bend, maximum tail bend, duration, tail-beat frequency, maximum tail curvature, and asymmetry, respectively). **C:** Comparison between asymmetry extracted directly from empirical tail movements and the corresponding fitted model values. Direct extraction of asymmetry from empirical data is more susceptible to noise because the signal has a lower dynamic range in asymmetric movements. **D-F:** Examples of compound movements composed of multiple concatenated component movements (197 concatenated movements in the dataset out of 7713). **D:** Component movements separated by a short temporal interval. **E:** Component movements concatenated without an intervening pause. Individual components could be identified either from the envelope or by abrupt changes in movement parameters, such as maximum tail curvature and tail-beat frequency. **F:** Component movements concatenated without an intervening pause. Individual components could be identified either from the envelope or by abrupt changes in maximum tail bend.

**Supplementary Movie 1: Fish movements and their PC coefficient dynamics.** Left: Recording of a fish performing four movements: one right turn, one left turn, and two forward swims, in which the tail oscillates symmetrically around the body axis. Upper right: Extracted tail movements (blue) and their reconstruction using the first three PCs (orange). Lower right: Dynamics of the three PC coefficients, shown from top and side views. The top view reveals rotation at an approximately constant radius, with each full rotation corresponding to one tail undulation. The side view reveals variation along the third dimension, with positive and negative values corresponding to rightward and leftward tail bends, respectively.
